# Flavins in Focus: Vitamin B_2_ as a Metabolic Keystone for Neuronal Repair in Regenerating Planarians

**DOI:** 10.64898/2026.10.12.758197

**Authors:** Martijn Heleven, Vincent Jaenen, Rik Maes, Josep F. Abril, Robbe Voglar, Charlotte Segnana, Tom Artois, Nelly D. Saenen, Francesc Cebrià, Karen Smeets

## Abstract

Regeneration imposes substantial metabolic demands to support the reconstruction of functional tissues. Essential vitamins are indispensable metabolic cofactors, yet whether their availability is spatially organized to support local metabolic demands within regenerative tissues remains largely unknown. Here, we identify previously undescribed flavin-rich cells, referred to as “flavinocytes”, that form an extensive network that spatially organizes vitamin B_2_ metabolism in the planarian *Schmidtea mediterranea*. Using endogenous flavin autofluorescence as a label-free *in vivo* metabolic readout, we show that this network consists of elongated cells that form longitudinal varicosity-containing strands closely associated with the central nervous system. RNAi-mediated disruption of riboflavin uptake and metabolism reduced flavin-associated autofluorescence and its spatial distribution within the animal, while affecting blastema formation, eye regeneration, and nervous system reconstruction. These findings demonstrate that riboflavin metabolism is required for successful regeneration. Single-cell transcriptomics, protein localization, and functional perturbations further suggest a reciprocal relationship between flavinocytes and the regenerating nervous system, pointing to coordinated neuro-metabolic interactions during tissue repair. Together, our findings demonstrate the importance of riboflavin metabolism for nervous system regeneration and provide new insight into the spatial organization of metabolism during tissue regeneration.

## 1. Introduction

The restoration of complex tissues after injury is an energetically and biosynthetically demanding process that requires the coordinated integration of signaling and metabolic programs^1–4^. Although the signaling pathways governing regeneration have been extensively characterized, far less is known about the dynamic metabolic environment in which these pathways operate, particularly how the availability of energy, reducing power, and essential cofactors shapes cellular behavior^5–7^. Increasing evidence indicates that these metabolic inputs not only support tissue regeneration but actively modulate signaling dynamics in space and time^3,8–11^.

A major challenge in regenerative biology is understanding how neighboring cell populations establish distinct metabolic identities despite sharing the same microenvironment^12–14^. Such metabolic specialization may arise not only from differences in pathway activity but also from the local availability, storage, mobilization, and utilization of metabolic resources, including energy substrates, biosynthetic precursors, and essential cofactors^13,15–17^. Resolving the spatial organization of these resources may provide a framework for understanding how distinct metabolic states are established and maintained across regenerating tissues. Among these resources, vitamins represent a critical yet comparatively understudied class of cofactors^18,19^.

Unlike glycogen, lipids, and other intracellularly stored metabolic resources, many vitamins cannot be synthesized *de novo*^20^. Moreover, compared to the readily buffered fat-soluble vitamins, the availability of water-soluble vitamins, including the B-vitamin family, is often constrained as they are minimally stored and must be continuously supplied or locally maintained to sustain cellular function^21,22^. This constraint may be particularly relevant in metabolically demanding tissues such as the nervous system^23,24^, raising a fundamental question: how do regenerating tissues, and the distinct cell populations within them, locally acquire and maintain supplies of these diffusible cofactors to meet the exceptional metabolic demands of tissue repair?

Among water-soluble vitamins, the redox-active molecule riboflavin (vitamin B_2)_ is of particular interest, as it cannot be synthesized by animals^25^. Consequently, animals rely entirely on dietary uptake, cellular transport, and intracellular conversion into the flavin cofactors flavin mononucleotide (FMN) and flavin adenine dinucleotide (FAD). These cofactors serve as essential electron carriers for hundreds of flavoproteins involved in mitochondrial respiration, ROS metabolism, antioxidant defense, neurotransmitter recycling, and numerous other redox-dependent processes^26–28^. At the same time, its limited storage capacity raises the possibility that tissues, particularly those with high metabolic demand, require specialized mechanisms to locally capture, retain, and utilize flavins^29,30^. Whether such mechanisms exist *in vivo* during tissue regeneration, and whether they involve dedicated cell populations, remains unknown.

Freshwater planarians provide a unique opportunity to unravel this question. Following injury, individuals of *Schmidtea mediterranea* can regenerate an entire organism from small body fragments, including a fully functional central nervous system, through the coordinated integration of stem cell activity, tissue patterning, and metabolic remodeling^31–33^. Their optical transparency and suitability for live imaging further allow direct visualization of fluorescent dyes or endogenous signals within intact, live animals, an approach not readily available in most regenerative systems^34,35^. Together, these features make planarians an ideal model for resolving how essential metabolic cofactors are distributed, retained, and utilized across tissues with distinct metabolic identities during regeneration.

Here, we investigated how riboflavin metabolism is spatially organized and functionally integrated across regenerating tissues. By combining live spectral imaging, RNAi-mediated knockdown, pharmacological perturbations, immunohistochemistry, and existing single-cell transcriptomic datasets, we identified a previously undescribed flavin-rich cellular network that spatially organizes vitamin B_2_ metabolism during regeneration. We demonstrate that this network is closely associated with the regenerating nervous system, depends on riboflavin uptake and metabolism, and links local flavin availability to neuronal regeneration and sensing.

## 2. Results

### 2.1 Spectral profiling identifies flavin-associated autofluorescence in live planarians

Biomolecules such as flavins, lipofuscin, porphyrins, and NADH exhibit unique autofluorescent (AF) spectra that provide a spectral fingerprint for their molecular identity^36,37^. To specifically identify flavin-associated AF, imaging at different excitation wavelengths followed by lambda spectral imaging was performed in live animals, as AF associated with endogenous metabolically active compounds rapidly diminishes upon chemical fixation (Fig. 1)^38^. Planarians were immobilized using linalool and mounted for live confocal imaging as described previously^39^ (Fig. 1A).

**Figure 1.**
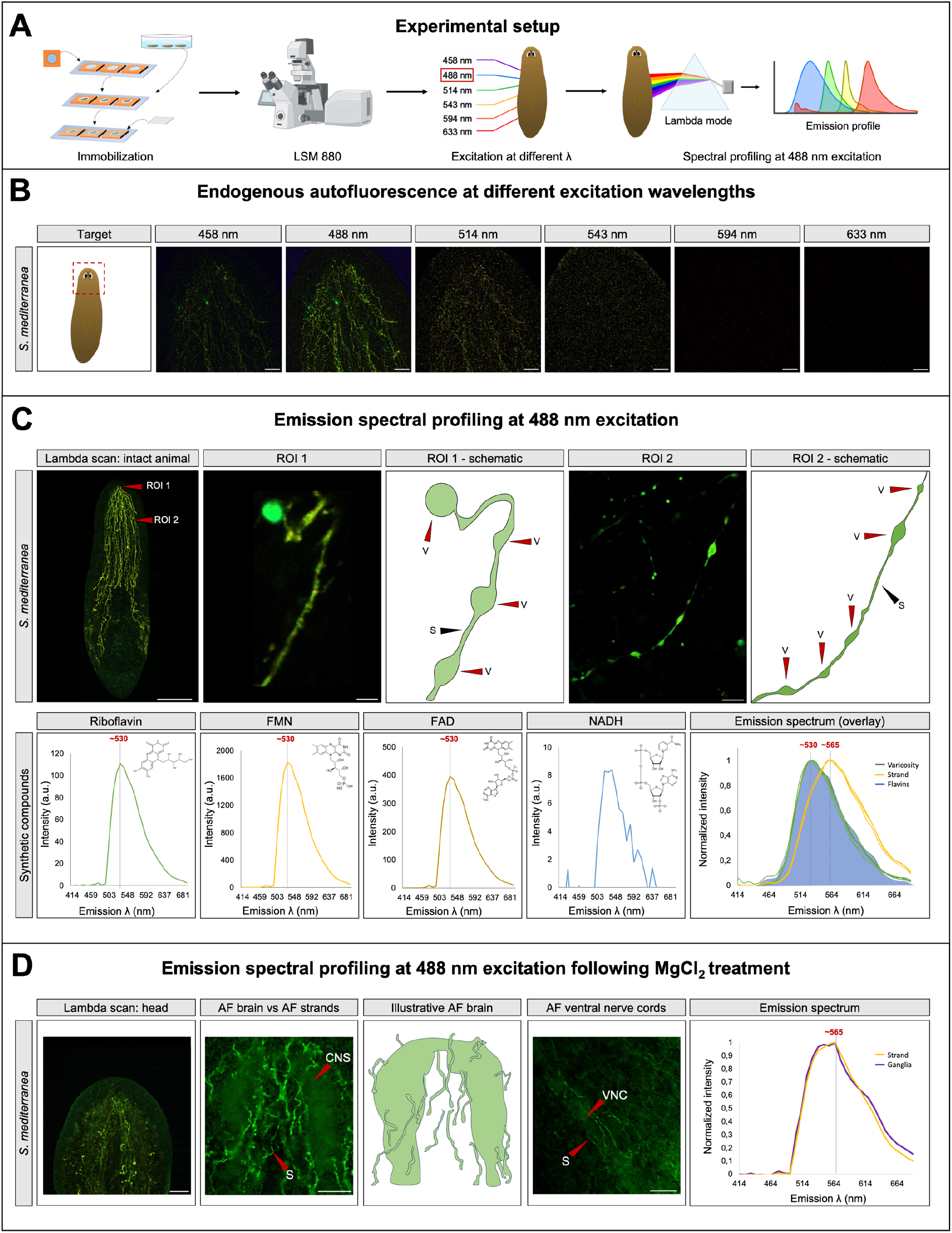
Spectral profiling identifies endogenous flavin autofluorescence in live *Schmidtea mediterranea*. **A)** Graphical representation of the experimental setup. Animals were immobilized and imaged using a Zeiss LSM 880 confocal microscope. Excitation at different wavelengths was performed using sequential laser lines (458–633 nm), followed by lambda-mode spectral imaging at 488 nm excitation to determine the emission profile of endogenous autofluorescent (AF) structures in comparison with synthetic compounds, including riboflavin, FMN, FAD, and NADH. Created in BioRender. Heleven, M. (2026) https://BioRender.com/1dxl2ep. **B)** Endogenous AF in the head region at different excitation wavelengths. Images show combined views of lambda-scan channels C1–C32 acquired at each excitation wavelength (458, 488, 514, 543, 594, and 633 nm; scale bars, 100 μm). **C)** Emission spectral profiling of endogenous AF at 488 nm excitation. Overview of the combined lambda-scan channels (C1–C32) in an intact live animal (scale bar, 500 μm), with higher-magnification views and corresponding schematics of structural details in ROI 1 and ROI 2 (scale bars, 20 μm), highlighting varicosity-like swellings (V) and strands (S). Emission spectra of synthetic riboflavin, FMN, FAD, and NADH are shown alongside the normalized emission spectra obtained from the corresponding ROIs, revealing emission maxima around ∼530 and ∼565 nm. Color legend: varicosity (green), strand (yellow), and synthetic flavins (blue). Lambda-scan imaging in control animals was performed in *n* ≥ 8 animals across at least three independent experiments. **D)** Emission spectral profiling at 488 nm excitation following MgCl_2 t_reatment. Combined view of lambda-scan channels C1–C32 showing AF in the head region following treatment with 0.5 M MgCl_2_ for 30 s (scale bar, 200 μm). Higher-magnification images and schematic show AF in the cephalic ganglia and adjacent strands, as well as AF in the ventral nerve cords (VNCs) (scale bars, 50 μm). Normalized emission spectra show a shared emission maximum around ∼565 nm in the ganglia and strands. Lambda-scan imaging following MgCl_2_ exposure was performed in *n* = 6 animals across two independent experiments.

Excitation at different wavelengths revealed that endogenous AF structures in the planarian head were most efficiently excited by blue light (458–514 nm), with maximal fluorescence observed at 488 nm. Longer excitation wavelengths (543–633 nm) generated only weak signals (Fig. 1B), supporting the use of 488 nm excitation for subsequent spectral analyses.

Lambda spectral imaging following 488 nm excitation of the head region identified two prominent endogenous emission peaks at ∼530 nm and ∼565 nm, corresponding to two spatially distinct cellular compartments (Fig. 1C). Comparison with synthetic flavins revealed that the endogenous ∼530 nm emission closely overlapped with the emission spectra of riboflavin, FMN, and FAD, while NADH was barely excited at 488 nm, supporting a flavin origin of the endogenous AF.

Structures such as the cephalic ganglia exhibited little to no detectable AF under standard *in vivo* imaging conditions compared with the strand-like structures. However, brief MgCl_2_ treatment, a commonly used method to induce planarian relaxation^40^, revealed clear AF within the ganglia and ventral nerve cords (Fig. 1D). Spectral profiling showed that these central nervous system structures exhibited emission profiles closely resembling those observed in the longitudinal AF strands, with a characteristic emission peak at ∼565 nm. This peak was approximately 35 nm red-shifted relative to the ∼530 nm peak associated with the punctate structures. Emission maxima at approximately 530 and 565 nm are characteristic of riboflavin and its derivatives and have been associated with free and protein-bound flavins, respectively^41–43^. Together, these findings demonstrate that endogenous AF in live planarians is spectrally consistent with endogenous flavins and reveal two spatially distinct flavin-associated autofluorescent components characterized by emission maxima at approximately 530 nm and 565 nm.

### 2.2 Flavin-associated autofluorescence identifies a distinct population of flavin-rich cells: flavinocytes

Flavin-associated AF identified a specific population of flavin-rich cells, hereafter termed flavinocytes (Fig. 2). Blue-light excitation at 488 nm revealed dense clusters within the head region that extended as longitudinal strands throughout the body. Spatially resolved spectral analysis demonstrated that the 530 nm emission predominated within varicosity-like structures, whereas the 565 nm emission was primarily associated with elongated longitudinal strand-like structures (Fig. 1C). Co-labeling with the superoxide probe to provide tissue-specific contrast demonstrated that the flavinocyte network was positioned ventral to the intestinal tract (Fig. 2A).

**Figure 2.**
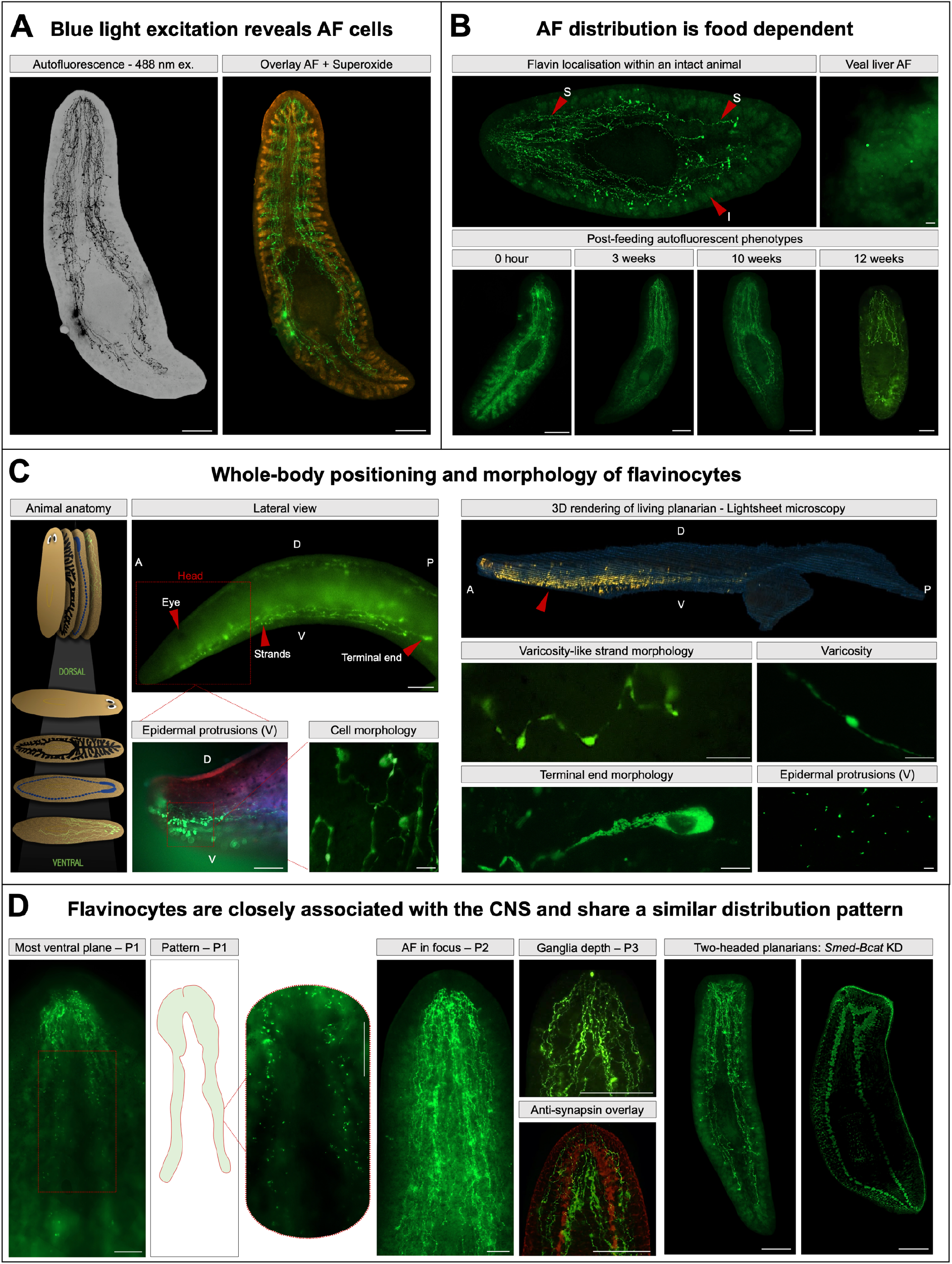
Flavin autofluorescence defines a distinct population of flavin-rich cells: flavinocytes. **A)** Blue light excitation reveals autofluorescent (AF) cells in live *Schmidtea mediterranea* (488 nm excitation). Whole-animal tilescans show AF at 488 nm excitation represented using an inverted LUT and an overlay of endogenous AF with a live superoxide-sensitive fluorescent probe (scale bars, 500 μm). **B)** AF distribution is food dependent. Distribution of flavin AF in an intact animal, highlighting strands (S) and intestine (I), and AF in veal liver (scale bar, 20 μm). Lower panels show post-feeding AF phenotypes at 0 h, 3 weeks, 10 weeks, and 12 weeks (scale bars, 500 μm). **C)** Whole-body positioning and morphology of flavinocytes. Anatomical overview, lateral view (scale bar, 100 μm), and three-dimensional light-sheet reconstruction illustrate the spatial organization and ventral localization of flavinocytes in live animals. Magnified views show epidermal protrusions (scale bar, 100 μm) and cell morphology (scale bar, 20 μm). High-resolution confocal images show varicosity-like strand morphology (scale bar, 20 μm), individual varicosities (scale bar, 10 μm), terminal end morphology (scale bar, 10 μm), and epidermal protrusions (scale bar, 5 μm). **D)** Flavinocytes are closely associated with the central nervous system (CNS) and share a similar distribution pattern. AF captured at different focal depths along the Z-axis is shown together with anti-synapsin immunostaining overlaid with flavinocytes observed *in vivo* within the same animal. Two-headed animals generated via *Smed-β-catenin* RNAi (*n* = 3) (scale bars, 200 μm).

The intensity and distribution of flavin-associated AF were strongly influenced by nutritional status (Fig. 2B). Well-fed animals displayed an extensive network of intensely fluorescent flavinocytes, while prominent AF was also observed throughout the intestinal tract following feeding. The intestinal fluorescence closely resembled the AF signal detected in veal liver, the flavin-rich food source used for feeding. During prolonged starvation, intestinal flavin-associated AF clearly declined. Nevertheless, the characteristic longitudinal flavinocyte network remained readily detectable, even after 12 weeks without food.

Whole-body imaging demonstrated that flavinocytes are localized at the ventral side of the animal, beneath the intestinal tract and in close proximity to the central nervous system (Fig. 2C). High-resolution confocal imaging and three-dimensional light-sheet microscopy further revealed that the densely clustered flavinocytes within the head extended fine cellular projections toward the ventral epidermis (Fig. 2C; Movie 1). Along the body axis, the longitudinal strands displayed regularly spaced varicosity-like swellings and terminated in enlarged distal endings, revealing a highly organized cellular network. Live imaging further revealed movement of autofluorescent material along the longitudinal strands, suggesting transport of cargo within the flavinocyte network (Movie 2). Additionally, Tubulin Tracker™ staining showed that flavinocyte-associated AF colocalized with tubulin-containing cellular processes, particularly within the head region and along the elongated longitudinal strands throughout the body (Fig. S1).

The anatomical organization of flavinocytes closely paralleled that of the central nervous system (Fig. 2D). Consistent with the shared 565 nm emission peak observed following MgCl_2_ treatment (Fig. 1D), flavinocytes occupied the same bilaterally symmetric ventral plane as the ventral nerve cords and converged anteriorly toward the cephalic ganglia. Within this plane, punctate AF structures became apparent throughout the ganglia and along the ventral nerve cords, further supporting their close anatomical association. Overlay of the endogenous AF signal with anti-synapsin immunostaining demonstrated that the flavinocyte network closely follows the organization of the central nervous system, consistent with the MgCl_2-_induced AF pattern observed in live animals (Fig. 1D). To further examine this relationship, *Smed-β-catenin* RNAi animals with duplicated heads were analyzed. Ectopic acquisition of head identity was accompanied by a corresponding duplication of the flavinocyte network, with flavinocytes clustering around both cephalic ganglia and extending along the duplicated ventral nerve cords. Together, these findings reveal a previously undescribed, bilaterally symmetric network of flavin-rich cells, whose organization in live animals closely mirrors the architecture of the planarian central nervous system previously defined in fixed specimens.

### 2.3 Riboflavin uptake and metabolism are broadly distributed and enriched in neural tissues

#### 2.3.1 Anatomical regions

Analysis of the published single-cell transcriptomic atlas (Fincher et al., 2018)^44^ revealed broad expression of riboflavin pathway genes across anatomical regions, with the highest transcript abundance in the head, prepharyngeal region, and trunk, and a lower expression posteriorly (Fig. 3A,B). *Smed-RFT1* and *Smed-RFT2* were detected throughout the body but at relatively low abundance, whereas *Smed-RFK*, *Smed-FADS*, and particularly *Smed-FMNr* showed substantially higher expression. Region-specific co-expression analysis identified the head as the anatomical region, forming the largest and most extensive riboflavin-related co-expression network (Fig. 3E).

**Figure 3.**
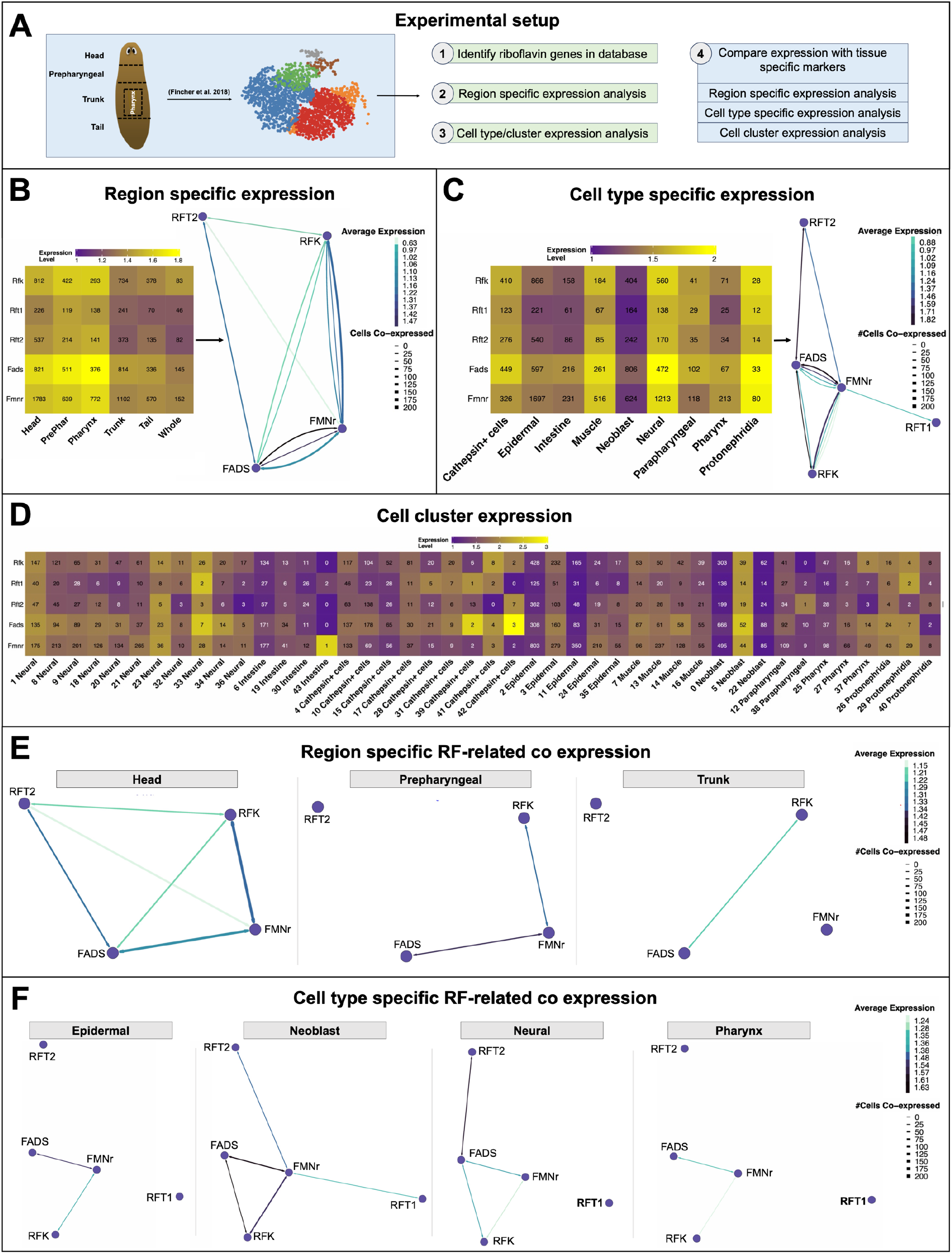
Molecular identity of riboflavin uptake and metabolism. **A)** Graphical representation of the experimental setup. Published single-cell RNA-sequencing data from Fincher et al. (2018) were analyzed to characterize the expression and co-expression of genes involved in riboflavin uptake and metabolism across anatomical regions, major cell types, and individual cell clusters. Created in BioRender. Heleven, M. (2026) https://BioRender.com/1dxl2ep. **B)** Region-specific expression of *Smed-RFT1*, *Smed-RFT2*, *Smed-RFK*, *Smed-FADS*, and *Smed-FMNr* across the head, prepharyngeal region, pharynx, trunk, tail, and whole animal. The heatmap shows expression levels of individual genes, while the network summarizes their relative expression and co-expression across anatomical regions. **C)** Cell-type-specific expression of riboflavin-related genes across major cell types. The heatmap shows expression levels of individual genes, while the accompanying network summarizes their relative expression and co-expression across cell types. **D)** Expression of riboflavin-related genes across individual transcriptionally defined cell clusters, illustrating the distribution of the riboflavin uptake and metabolic machinery at higher resolution. **E)** Region-specific co-expression networks of riboflavin-related genes in the head, prepharyngeal region, and trunk. **F)** Cell-type-specific co-expression networks of riboflavin-related genes in epidermal, neoblast, neural, and pharyngeal cell types. In all the co-expression network representations, edge color represents the average gene expression while edge thickness represents the number of co-expressing cells.

#### 2.3.2 Cell types and clusters

Across major cell types and transcriptionally defined cell clusters, riboflavin pathway genes displayed a broad and heterogeneous expression pattern (Fig. 3C,D). Neural, cathepsin+, and muscle populations exhibited the highest transcript abundance, whereas epidermal cells contained the greatest numbers of expressing cells despite lower expression levels. Cell type-specific co-expression analysis further identified neural and neoblast populations as the primary sites of coordinated riboflavin pathway gene expression (Fig. 3F). Analysis of additional published single-cell transcriptomic atlases^45,46^ further supported the broad distribution of riboflavin pathway genes across multiple cell types (Fig. S2).

Comparison with established neuronal subtype markers demonstrated broad expression of riboflavin pathway genes across aminergic, cholinergic, GABAergic, glutamatergic, glycinergic, panneuronal, and peptidergic neuronal populations (Fig. S3). Consistent with previous analyses, *Smed-FMNr* showed the highest transcript abundance, whereas *Smed-RFT1* and *Smed-RFT2* were expressed at lower levels. No exclusive association with a single neuronal subtype was observed. Co-expression network analysis further demonstrated coordinated co-expression of riboflavin-related genes across all major neuronal populations, with the strongest networks observed for panneuronal, peptidergic, cholinergic, and glycinergic markers (Fig. 4D).

**Figure 4.**
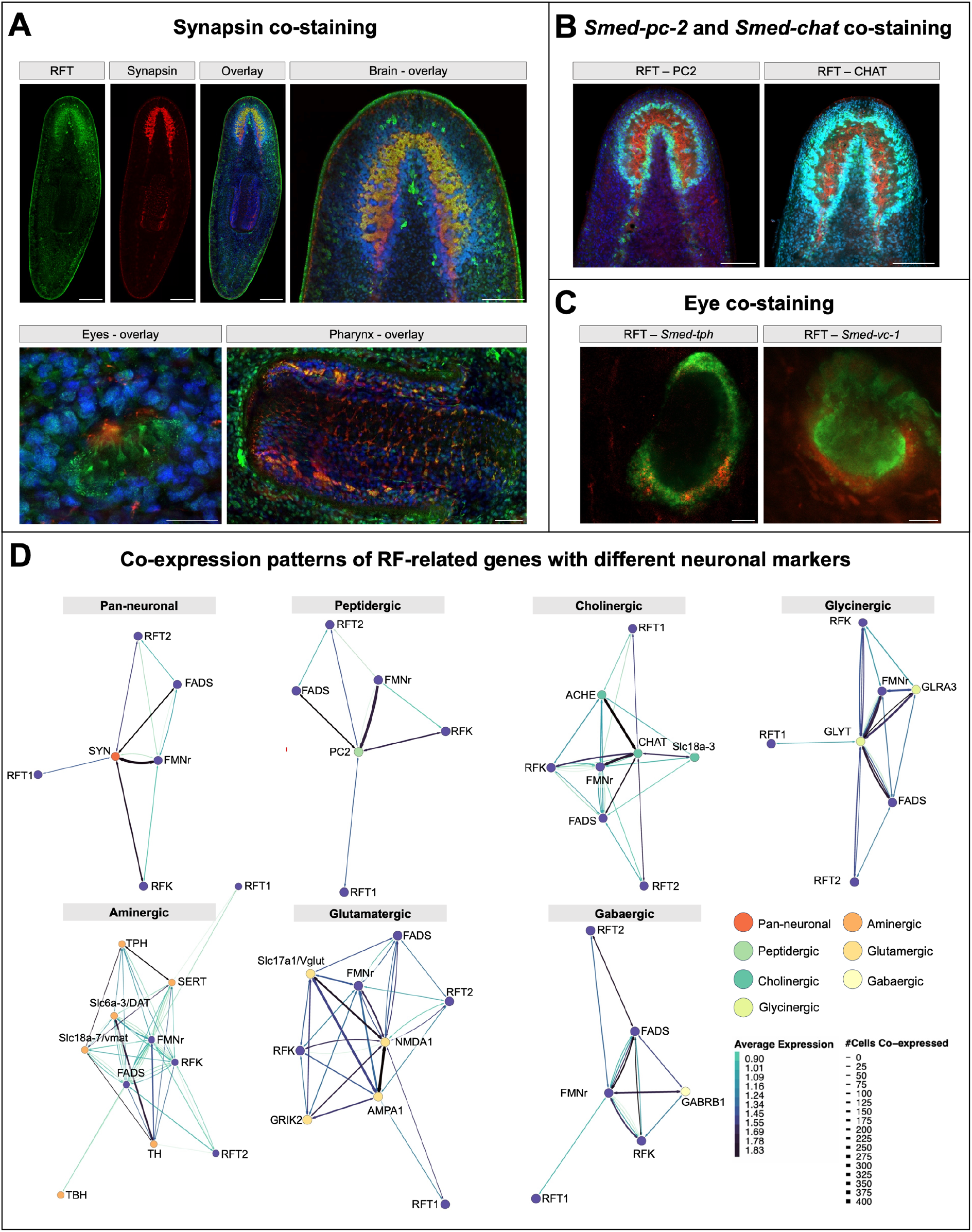
Riboflavin transporter colocalization with neuronal markers. **A)** Whole-mount immunostaining of riboflavin transporter (RFT; green), synapsin (red), and Hoechst (blue). Single-channel images, merged overview, and zoomed-in insets of the brain are shown (scale bars, 100 μm), alongside images of the eye (scale bar, 20 μm) and pharynx (scale bar, 50 μm). **B)** Whole-mount immunostaining of RFT combined with FISH for the neuronal markers *Smed-pc2* and *Smed-chat*. Color code: RFT (red), neuronal markers (green), Hoechst (blue) (scales, 100 μm). **C)** Whole-mount immunostaining of RFT combined with FISH for the eye-associated markers *Smed-tph* and *Smed-vc-1*. Color code: RFT (red), eye-associated markers (green), Hoechst (blue) (scales, 10 μm). **D)** Co-expression networks of riboflavin-related genes with neuronal subtype markers using the Fincher et al. (2018) single-cell RNA-sequencing atlas. Networks are shown for pan-neuronal, peptidergic, cholinergic, glycinergic, aminergic, glutamatergic, and GABAergic neuronal populations. Edge color represents the average gene expression while edge thickness represents the number of co-expressing cells.

RFT immunostaining confirmed the neural enrichment predicted by the transcriptomic analyses, showing strong localization within the cephalic ganglia, ventral nerve cords, and pharyngeal nervous system, with extensive overlap with Synapsin (Fig. 4A). High-resolution imaging further demonstrated RFT-positive structures surrounding the cephalic ganglia, photoreceptor neurons, and pharyngeal nerve net. Co-staining with markers for peptidergic (PC2), cholinergic (CHAT), serotonergic (TPH), and visual (VC1) neurons demonstrated close association with multiple neuronal subtypes (Fig. 4B,C).

Although neural populations exhibited the strongest coordinated expression, riboflavin pathway genes were also detected in cathepsin^+^ (glial/parenchymal) populations, with *Smed-RFT2* showing enrichment in specific cathepsin^+^ clusters, whereas *Smed-RFK*, *Smed-FADS*, and *Smed-FMNr* were distributed more broadly across multiple clusters (Fig. S4).

The flavinocyte network was anatomically distinct from the planarian excretory system (Fig. S5). Fluorescent *in situ* hybridization for the excretory marker *Smed-inx10* showed a protonephridial pattern that remained spatially distinct from flavin-associated AF (Fig. S5B). Moreover, *in vivo* labeling of the excretory system with fluorescent dextran (500-kDa) revealed a characteristic branched tubular morphology that remained anatomically distinct from the flavinocyte network and was positioned more dorsally^47^ (Fig. S5C).

### 2.4 Disruption of riboflavin uptake and metabolism impairs (neuro-)regeneration

#### 2.4.1 RF-related interference disrupts the spatial organization of flavin-associated AF during regeneration

Interference with riboflavin uptake and metabolism altered both the intensity and spatial distribution of flavin-associated AF during regeneration (Fig. 5).

**Figure 5.**
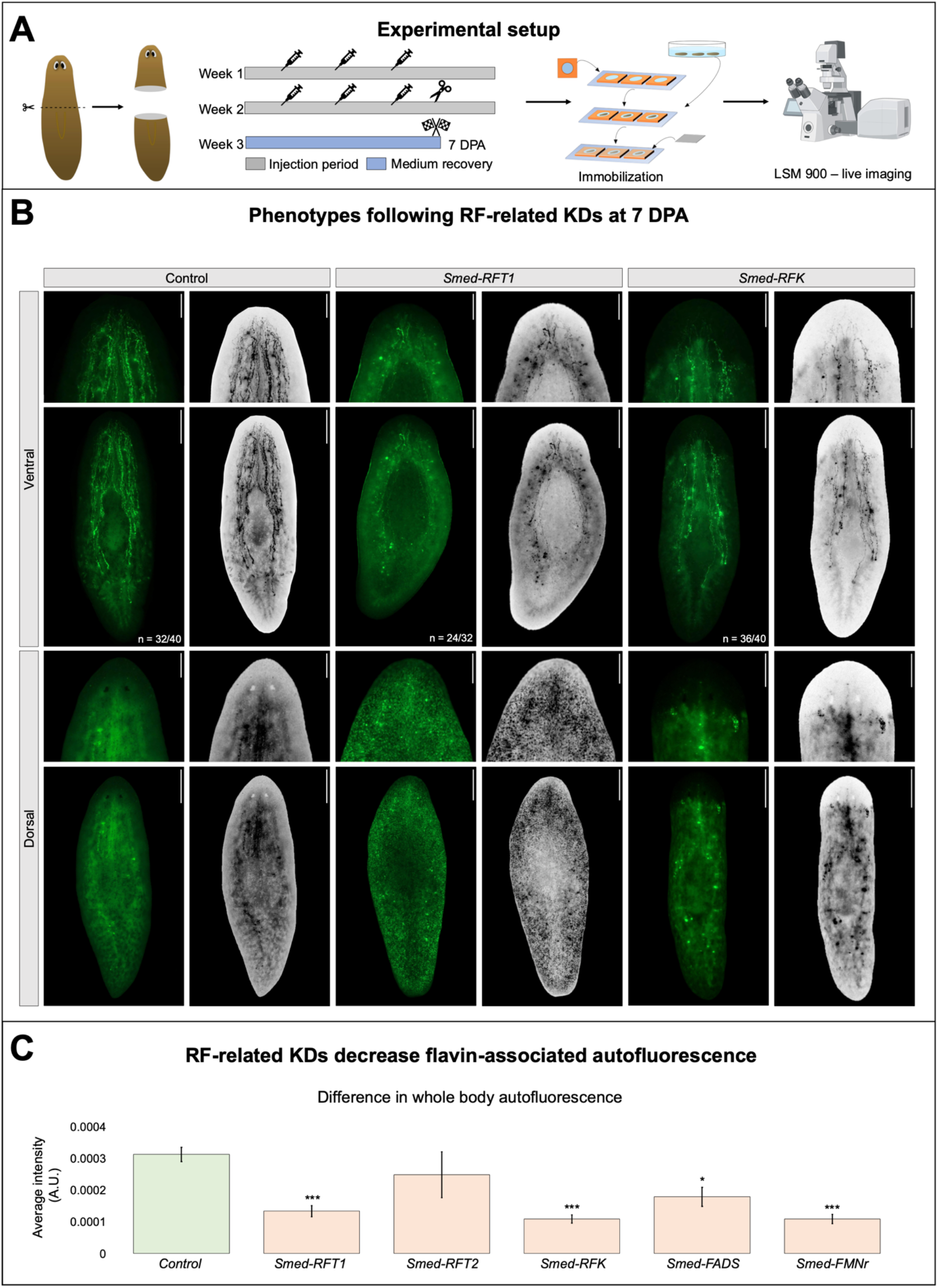
Riboflavin-related gene knockdown alters flavin autofluorescence in regenerating tails. **A)** Graphical representation of the experimental setup. Animals were subjected to two rounds of RNA interference (RNAi), each consisting of three injection days, targeting riboflavin-related genes. Following RNAi, animals were amputated prior to the pharynx and allowed to regenerate. At 7 days post-amputation (7 DPA), planarians were immobilized for live imaging using a Zeiss LSM900 confocal microscope. Created in BioRender. Heleven, M. (2026) https://BioRender.com/1dxl2ep. **B)** Representative live images of a selected subset of RF-related knockdowns, including RNAi of *Smed-RFT1* and *Smed-RFK* are shown. Whole-animal views (scale bars, 500 µm) and zoomed-in insets (scale bars, 250 µm) of the head region are shown from both dorsal and ventral sides. Autofluorescence (AF) display intensity was adjusted independently between images to visualize differences in AF distribution; images should therefore not be used for direct intensity comparison. The phenotypes shown were observed across at least three independent experiments. **C)** Quantification of raw whole-body flavin-associated AF in control and RNAi-treated animals, normalized against the body area. *n* = 16 animals per group across two independent experiments. Representative phenotypes corresponding to the other flavin-related genes are summarized in Fig. S6. The indicated *n* represents the total number of animals showing the respective phenotype. Statistical significance is indicated by: ∗, p < 0.05; ∗∗, p < 0.01; ∗∗∗, p < 0.001.

Control animals regenerated a flavin-associated AF pattern closely resembling that of intact animals, consisting of a dense head cluster of AF-positive flavinocytes connected to longitudinal AF-positive strands extending throughout the body. Accordingly, flavin-associated AF was distributed throughout the head region and was readily detected from both dorsal and ventral perspectives (Fig. 5B; Fig. S6B).

Following *Smed-RFT1* RNAi, flavin-associated AF accumulated predominantly within the dorsal epidermis, while AF associated with the head cluster and longitudinal strands was markedly reduced. Consequently, flavin-associated AF became largely restricted to a small head-associated cluster and a limited number of longitudinal strand-like structures. A similar, although less pronounced, phenotype was observed following *Smed-RFT2* RNAi (Fig. S6B).

Interference with the downstream metabolic enzymes resulted in progressively more severe alterations in flavin-associated AF. *Smed-RFK*, *Smed-FADS*, and *Smed-FMNr* RNAi caused a marked reduction in whole-body flavin-associated AF, accompanied by a substantial loss of AF associated with the head region, longitudinal strands, and regenerating blastema. These effects were most pronounced following *Smed-RFK* and *Smed-FMNr* knockdown (Fig. S6B).

Quantification of whole-body flavin-associated AF confirmed a significant reduction in AF intensity following interference with riboflavin uptake and metabolism (Fig. 5C).

#### 2.4.2 RF-related interference impairs blastema and eye formation

RNAi-mediated disruption of riboflavin uptake and metabolism resulted in distinct regeneration phenotypes (Fig. 6A,B). Knockdown of *Smed-RFK* and *Smed-FMNr* significantly reduced blastema size. In contrast, *Smed-RFT1*, *Smed-RFT2*, and *Smed-FADS* did not produce clear defects in blastema size compared with control animals at 7 DPA (Fig. 6C,D).

**Figure 6.**
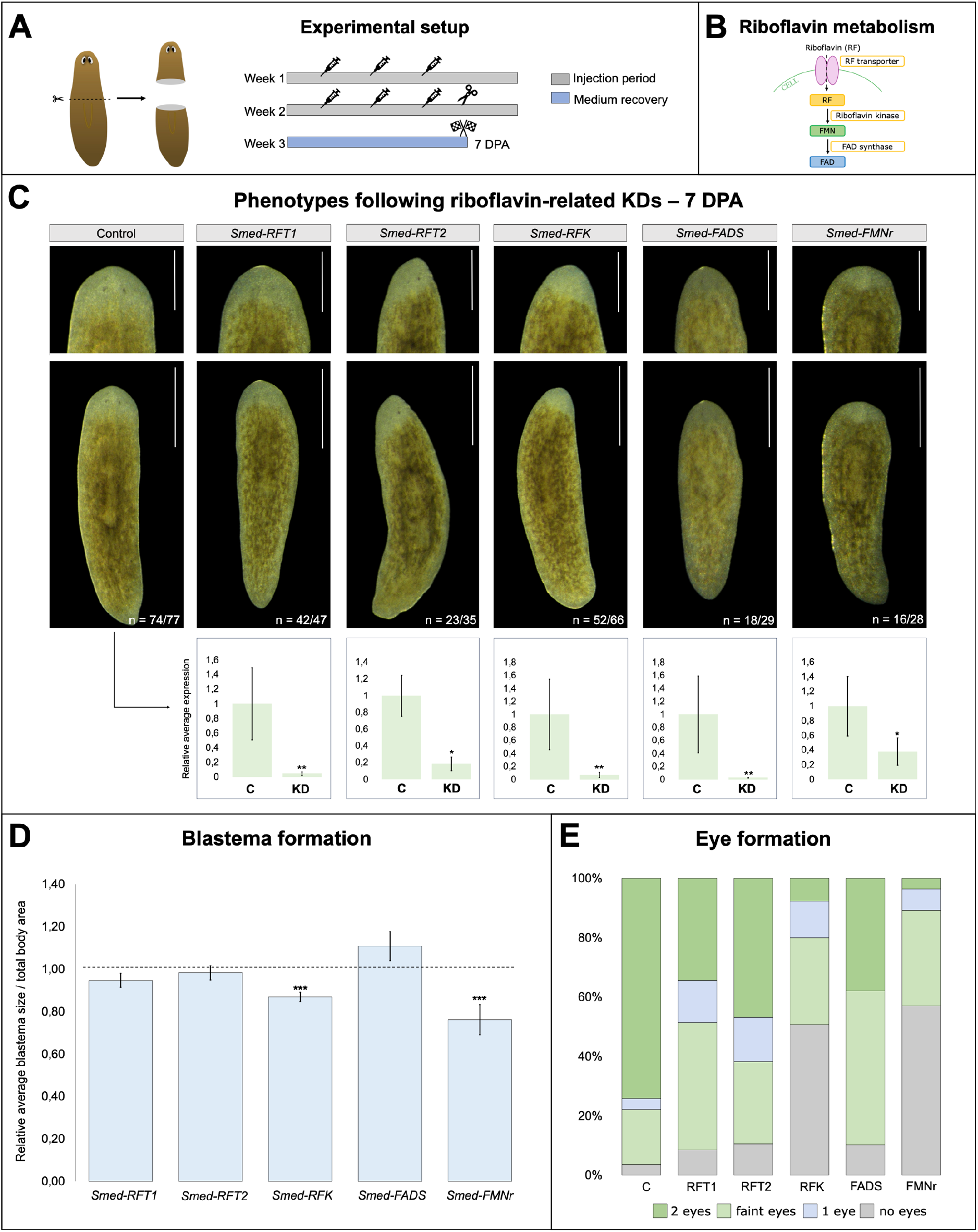
Interfering with riboflavin uptake and metabolism affects head regeneration. **A)** Graphical representation of the experimental setup. Animals were subjected to two rounds of RNA interference (RNAi), each consisting of three injection days, targeting riboflavin-related genes. Following RNAi, animals were amputated prior to the pharynx and allowed to regenerate. At 7 days post-amputation (7 DPA), phenotypic assessment was performed. **B)** Schematic overview of the riboflavin uptake and metabolic pathway, including riboflavin transport (RFT), phosphorylation to flavin mononucleotide (FMN) via riboflavin kinase (RFK), and conversion to flavin adenine dinucleotide (FAD) via FAD synthase (FADS). **C)** Representative regenerating animals following control RNAi or RNAi of *Smed-RFT1*, *Smed-RFT2*, *Smed-RFK*, *Smed-FADS*, and *Smed-FMNr* at 7 DPA. Whole-animal views (scale bars, 500 µm) and zoomed-in insets (scale bars, 250 µm) of the regenerated head region are shown. Graphs below the representative phenotypes show knockdown efficiency validated by RT-qPCR. **D)** Quantification of regenerated blastema area normalized to total body area following RNAi and expressed relative to the corresponding control. The dashed line indicates the normalized control value of 1. **E)** Quantification of eye phenotypes shown in C. Eye phenotypes were scored according to the presence of two eyes, two faint eyes, one eye, or no eyes. Data were acquired across two to five independent experiments. The indicated *n* represents the total number of animals showing the respective phenotype. Statistical significance is indicated by: *, *p* < 0.05; **, *p* < 0.01; ***, *p* < 0.001.

The severity of impaired blastema growth was broadly reflected in the extent of eye regeneration defects at 7 DPA (Fig. 6E). Most control animals regenerated two normally pigmented eyes, whereas perturbation of riboflavin metabolism increased the frequency of eye regeneration defects to varying degrees. The most pronounced phenotypes were observed following *Smed-RFK* and *Smed-FMNr* knockdown, with many animals exhibiting absent (RFK, 50%; FMNr, 57%) or faintly pigmented eyes (RFK, 29%; FMNr, 32%). In contrast, *Smed-RFT1*, *Smed-RFT2*, and *Smed-FADS* knockdown produced comparatively mild defects.

#### 2.4.3 RF-related interference impairs neuroregeneration

Whole-mount anti-synapsin staining at 7 DPA revealed a significant reduction in regenerated brain area following RNAi-mediated knockdown of all riboflavin pathway genes, with the strongest defects observed after *Smed-RFK* and *Smed-FMNr* RNAi (Fig. 7B,C).

**Figure 7.**
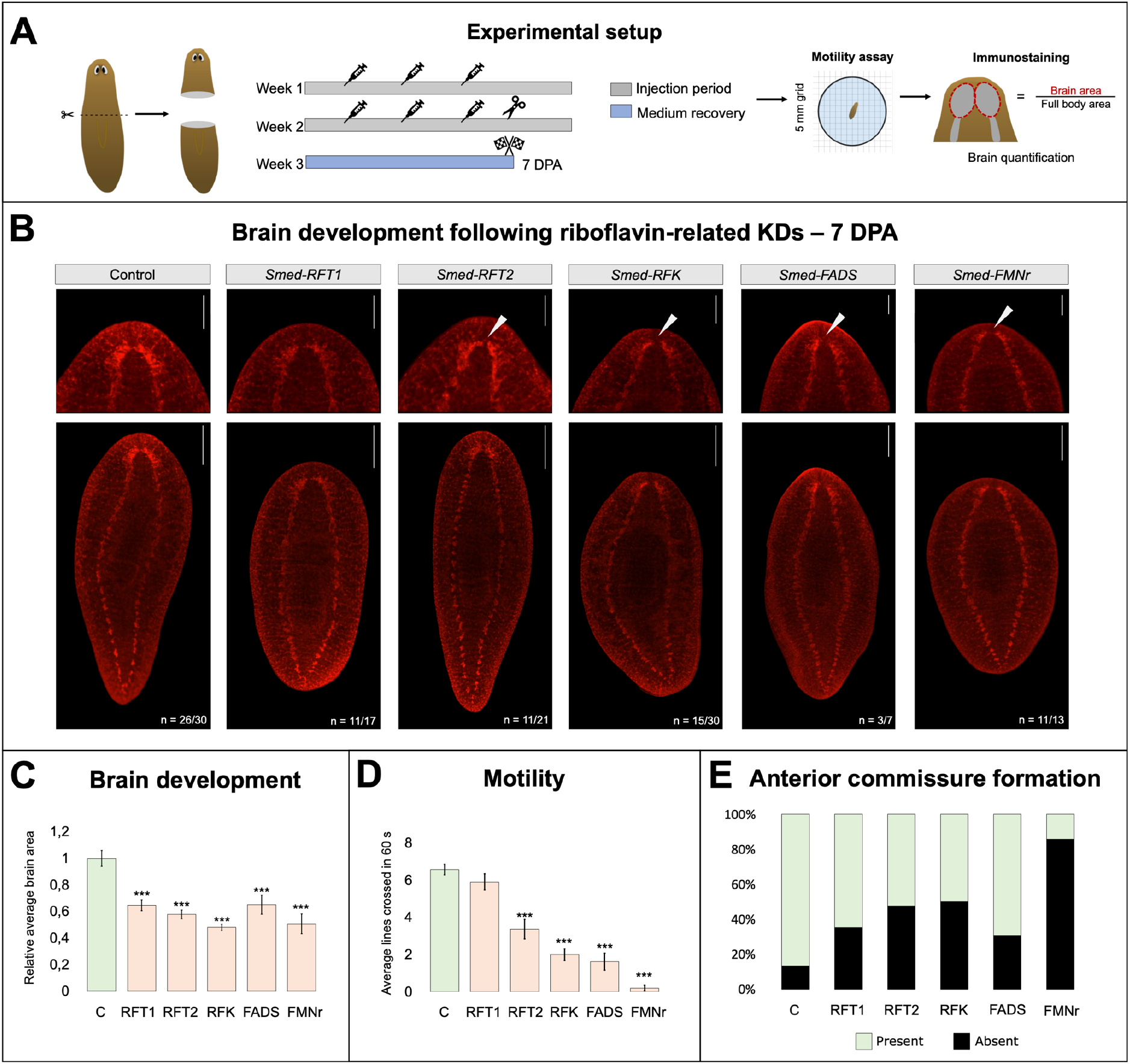
Interfering with riboflavin uptake and metabolism affects nervous system regeneration. **A)** Graphical representation of the experimental setup. Animals were subjected to two rounds of RNA interference (RNAi), each consisting of three injection days, targeting riboflavin-related genes. Following RNAi, animals were amputated prior to the pharynx and allowed to regenerate. At 7 days post-amputation (7 DPA), motility assessment and anti-synapsin immunostaining were performed to quantify brain development. **B)** Representative anti-synapsin immunostaining of the regenerated brain following control RNAi or RNAi of *Smed-RFT1*, *Smed-RFT2*, *Smed-RFK*, *Smed-FADS*, and *Smed-FMNr* at 7 DPA. Whole-animal views (scale bars, 200 µm) and zoomed-in insets (scale bars, 100 µm) of the regenerated head region are shown. White arrows indicate the absence of the anterior commissure. **C)** Quantification of regenerated brain area following RNAi. Brain area is normalized to the total body area and expressed relative to the corresponding control group. **D)** Quantification of motility following RNAi. The average number of lines crossed during a 60 s interval is shown. **E)** Quantification of anterior commissure formation. Animals were categorized according to the presence or absence of the anterior commissure. Data were acquired across two independent experiments. The indicated *n* represents the total number of animals showing the respective phenotype. Statistical significance is indicated by: *, *p* < 0.05; **, *p* < 0.01; ***, *p* < 0.001.

These morphological defects were accompanied by impaired locomotor behavior. Animals subjected to *Smed-RFT2*, *Smed-RFK*, *Smed-FADS*, and *Smed-FMNr* RNAi showed a pronounced reduction in motility, with *Smed-FMNr* knockdown resulting in an almost complete loss of locomotion, whereas *Smed-RFT1* RNAi had no effect (Fig. 7D). Consistent with this, formation of the anterior commissure was also disrupted following RNAi, with no commissure formed most frequently after *Smed-FMNr* (86%), *Smed-RFK* (50%), and *Smed-RFT2* (48%) RNAi (Fig. 7B,E).

### 2.5 Neuronal signaling regulates the spatial organization of flavin-associated autofluorescence during regeneration

Genetic and pharmacological interference with neuronal signaling similarly altered the characteristic flavin-associated AF pattern during regeneration at 7 DPA (Fig. S7–S8).

Knockdown of the peptidergic marker *Smed-pc2* affected flavinocyte-associated AF, characterized by reduced AF within the head-associated flavinocyte cluster and along the longitudinal flavinocyte strands. Similarly, inhibition of cholinergic (*Smed-chat*), serotonergic

(*Smed-tph*), dopaminergic (*Smed-th),* and octopaminergic (*Smed-tbh*) pathways altered the characteristic flavinocyte-associated AF pattern, although the severity differed between neuronal populations (Fig. S7B). RNAi-mediated knockdown of the serotonergic neuronal specification and maintenance factors *Smed-lhx1/5-1* and *Smed-pitx*^48^ also resulted in severe disruption of the characteristic flavinocyte-associated AF pattern. In contrast, *Smed-gad* RNAi produced no visibly detectable alterations in flavinocyte-associated AF.

To independently validate these RNAi-induced observations, regenerating animals were continuously exposed to pharmacological inhibitors targeting neuronal activity and neurotransmission (Fig. S8). Inhibition of voltage-gated sodium channels with tetrodotoxin or mitochondrial respiration with rotenone impaired spatial AF distribution, resulting in little to no flavin-associated AF associated with the head cluster and longitudinal strands. Likewise, pharmacological interference with dopaminergic signaling using 3-iodo-L-tyrosine or 6-hydroxydopamine significantly reduced flavinocyte-associated AF, resulting in little to no AF associated with the head cluster and markedly reduced AF along the longitudinal strands. Similar alterations in flavinocyte-associated AF were observed following disruption of serotonergic signaling with ondansetron or fluoxetine and cholinergic neurotransmission with atropine or hexamethonium. Interestingly, serotonergic signaling perturbed animals also showed a shift of flavin-associated AF toward the intestinal tract compared to the control, a phenotype also observed consistently following *Smed-pitx* RNAi. Although the severity of the phenotypes varied between compounds, all treatments altered the characteristic flavinocyte-associated AF pattern during regeneration.

### 2.6 Mature flavinocytes dynamically respond to environmental stress

Flavinocyte-associated AF rapidly changed in response to acute environmental and physiological perturbations (Fig. 8).

**Figure 8.**
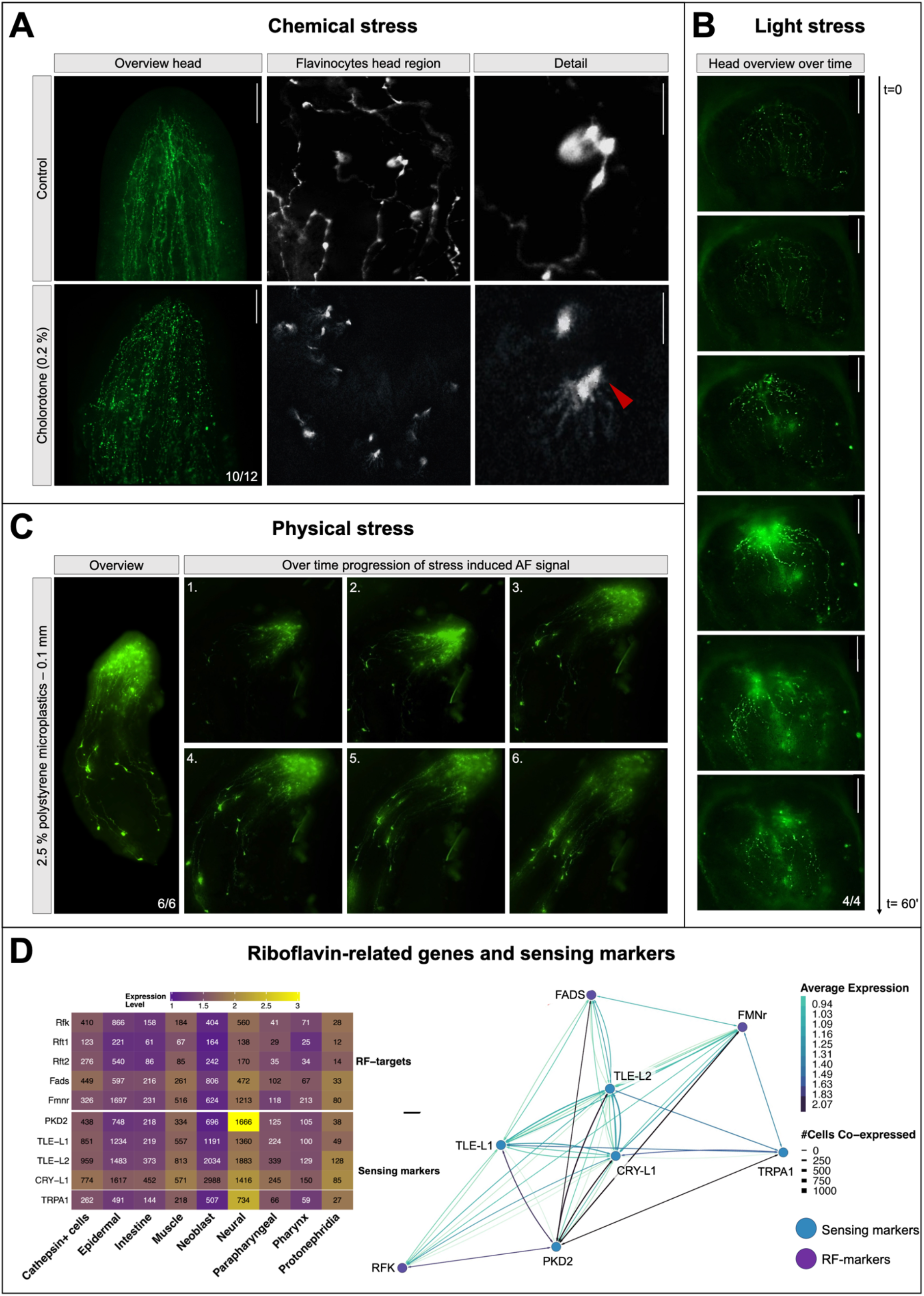
Stress reactions of flavinocytes under different stimuli. Summary of the phenotypes observed following chemical, physical, and light stimulation. **A)** Representative live images of flavinocyte-associated autofluorescence (AF) in control animals compared to animals exposed to 0.2% chlorotone for 10 min. Overview images of the head region (scale bars, 200 µm), higher-magnification images of flavinocytes, and zoomed-in insets (scale bars, 20 µm) are shown. **B)** Time-lapse imaging of flavinocyte-associated AF during continuous blue light excitation over a 60 min imaging period. **C)** Representative time-lapse images of flavinocyte-associated AF following physical stimulation using a solution of 2.5% polystyrene particles (0.1 mm). Time-lapse images illustrate the progression of the endogenous AF signal following stimulation. **D)** Expression heatmap and co-expression network of riboflavin-related genes and well-established sensory-associated markers derived from the *Schmidtea mediterranea* single-cell RNA-sequencing atlas. The corresponding co-expression network is shown on the right. Line color represents the average gene expression while line thickness represents the number of co-expressing cells. The indicated *n* represents the total number of animals showing the respective phenotype.

Brief exposure (10 min) to chlorotone, an anesthetic commonly used for planarian immobilization^49^, rapidly altered the characteristic flavinocyte-associated AF pattern (Fig. 8A). In control animals, flavinocyte-associated AF showed a continuous network of longitudinal strands interconnected by prominent varicosity-like swellings that converged into a dense cluster within the head region. In contrast, chlorotone-treated animals showed a clear reduction in AF associated with the interconnecting strands, whereas the enlarged varicosity-like swellings remained readily detectable. High-resolution imaging confirmed fragmentation of strand-associated AF, suggesting that chlorotone primarily disrupts the spatial organization of flavin-associated AF rather than abolishing the signal entirely (Fig. 8A).

To determine whether flavin-associated AF dynamically responds to light stimuli, intact animals were continuously exposed to blue-light excitation for up to 60 min during live imaging (Fig. 8B). This induced a rapid increase in flavin-associated AF that propagated throughout the flavinocyte network.

Next, physical stimulation was induced by exposing animals to 0.1 mm polystyrene microplastics. Compared with untreated controls, microplastic-exposed animals exhibited a rapid increase in flavin-associated AF within minutes, characterized by enhanced intracellular AF accumulation, particularly within the head region and at terminal flavinocyte endings throughout the body (Fig. 8C).

Because flavinocyte-associated AF was dynamically altered by chemical, physical, and light stimulation, we investigated whether riboflavin pathway genes were transcriptionally associated with sensory markers. Comparison with established sensory markers revealed coordinated expression of riboflavin pathway genes within neural tissue, with particularly strong co-expression observed with the sensory marker *Smed-pkd2*, together with extensive transcriptional associations with *Smed-tle-l1*, *Smed-tle-l2*, *Smed-cry-l1*, and *Smed-trpa1* (Fig. 8D). Since *Smed-pkd2* has recently been shown to mark sensory neurons involved in mechanosensation and food detection in planarians, these findings further support a potential association between riboflavin metabolism and sensory neuronal function.

### 2.7 Flavinocyte-associated autofluorescence networks are detectable in multiple flatworm species

The flavinocyte-associated AF network, morphologically, spectrally, and functionally characterized here in asexual strains of *Schmidtea mediterranea*, was also observed in the asexual planarian *Dugesia japonica*, displaying a comparable spatial organization (Fig. S9). In contrast, comparable flavinocyte-like structures were not detected in *Macrostomum lignano*. Together, these observations indicate that the flavinocyte-associated AF network is present across multiple planarian species but is not universally detectable among flatworms.

## 3. Discussion

Essential vitamins are among the most ubiquitous metabolic cofactors across living systems, yet whether their availability is organized to support local metabolic demands within tissues, particularly at the whole-organism level, remains largely unexplored^19^. Here, we identify a previously undescribed flavin-rich cellular network that spatially organizes vitamin B_2_ metabolism during regeneration, suggesting that the spatial regulation of cofactor availability represents an additional layer of metabolic organization during tissue repair (Fig. 2). The existence of a dedicated flavin-rich cellular network suggests that the availability of water-soluble vitamins may not rely solely on systemic delivery and passive diffusion but may instead be regulated locally to meet the distinct metabolic demands of neighboring tissues or different cell types. Such local organization is particularly advantageous in highly regenerative tissues and organisms, where extensive cellular plasticity creates dynamic and spatially heterogeneous demands for metabolic cofactors^4,12^.

### 3.1 Blue-light-induced autofluorescence reflects endogenous riboflavin metabolism

Spatial organization of metabolism has been observed in previous studies across multiple biological scales, including metabolic zonation within vertebrate organs such as the liver and small intestine^3,13^, and regional or cell type-specific differences in flavin distribution in invertebrates such as earthworms^82^. Here, we extend these observations by demonstrating that vitamin B_2_ metabolism is organized through a previously undescribed flavin-rich cellular network spanning the regenerating planarian *Schmidtea mediterranea* (Fig. 2). We established that the green autofluorescence (AF) in this planarian, observed at 488 nm excitation, originates predominantly from flavins, with emission peaks matching those reported across diverse organisms and cell types (Fig. 1B,C)^41,50–55^. Consistent with previous spectroscopic studies, the ∼530 nm emission corresponds predominantly to free flavins, whereas the ∼565 nm peak likely reflects protein-bound flavins within flavoproteins^41–43^. Knockdown of riboflavin (RF) uptake and metabolism decreased overall organismal AF, further highlighting the flavin nature of this AF signal (Fig. 5; Fig. S6). Together, these findings establish endogenous flavin AF as a label-free readout of vitamin B2 metabolism in live planarians.

Flavin AF has long been exploited as a metabolic biomarker across a wide range of biological systems, including cancer, stem cells, immune cells, and other metabolically active cell populations^56–58^, where it serves as a proxy for cellular metabolism and redox state^51,53,59–61^. The identification of a dedicated flavin-rich cellular network (Fig. 1; Fig. 2) in planarians, therefore, indicates that specialized cellular architectures with endogenous flavin AF may coordinate local cofactor availability during regeneration.

### 3.2 Riboflavin metabolism is required for neuroregeneration

Because riboflavin is obtained primarily through dietary uptake prior to being converted into its active cofactors FMN and FAD, tissue-specific uptake, storage, and metabolism are critical determinants of local flavin availability^62^. Consistent with this, RNAi-mediated disruption of riboflavin transport and metabolism impaired regeneration, reducing blastema formation, eye development, and whole-animal flavin AF (Fig. 5; Fig. 6; Fig. S6). Phenotypic severity differed between pathway steps, with riboflavin kinase (*Smed-RFK*) producing the strongest defects (Fig. 6).

The most pronounced consequences of disrupting riboflavin metabolism were observed during nervous system regeneration (Fig. 7). Perturbation of riboflavin uptake or metabolism impaired brain regeneration, reduced anterior commissure formation, disrupted eye regeneration, and compromised locomotor behavior (Fig. 7C,D,E). Neurons are highly dependent on flavoprotein activity, as FMN and FAD serve as cofactors for mitochondrial enzymes of oxidative phosphorylation, antioxidant defense and neurotransmitter metabolism^63–65^.

This dependence appears conserved across metazoans. In humans, pathogenic variants in the riboflavin transporters SLC52A2 and SLC52A3 cause riboflavin transporter deficiency, a progressive neurodegenerative disorder characterized by motor and sensory neuropathies^23,66,67^. Likewise, dietary or genetic riboflavin deficiency in both vertebrate and invertebrate models, including *D. rerio* and *C. elegans*, produces mitochondrial dysfunction accompanied by neurodegeneration and altered cholinergic transmission^68,69^. Our findings extend these observations by demonstrating that riboflavin metabolism is not only required for neuronal maintenance and function, but also plays an essential role during nervous system regeneration following injury (Fig. 7; Fig. 8).

Consistent with this role in nervous system regeneration, riboflavin transport and metabolic genes are expressed across multiple cell populations but relatively enriched within neural lineages (Fig. 3; Fig. 4D; Fig. S2; Fig. S3; Fig. S4), while immunolocalization confirmed the presence of riboflavin transporters across the central nervous system (Fig. 4). Specifically, RFT immunoreactivity extensively overlapped with synapsin and was closely associated with peptidergic, cholinergic, and eye-associated neuronal populations (Fig. 4A-C), indicating that flavin uptake machinery is broadly distributed across neuronal subtypes rather than restricted to a specific lineage. Consistent with this, enhancing endogenous AF using MgCl_2_ revealed the protein-bound flavin emission peak (∼565 nm) within the cephalic ganglia and ventral nerve cords, although these structures exhibited relatively little intrinsic AF under basal imaging conditions (Fig. 1D; Fig. 2D). This observation suggests that the planarian central nervous system (CNS) contains a substantial pool of protein-bound flavins, which generally exhibit lower fluorescence quantum yields than free flavins^70^. Although this interpretation requires direct biochemical validation, it suggests that flavin metabolism is closely integrated with the planarian CNS.

### 3.3 Flavinocytes: flavin-enriched cells closely associated with the nervous system

Flavin AF has most commonly been described as a diffuse, cell-intrinsic signal reflecting mitochondrial metabolic state, with elevated fluorescence levels resembling highly metabolic tissues^60,61^. In planarians, high levels of flavin-associated AF were predominantly organized into a discrete and anatomically stereotyped cellular network spectrally characterized by two prominent emission peaks at 530 and 565 nm (Fig. 1C). We refer to these flavin-enriched cells as “flavinocytes”. To our knowledge, neither the cellular morphology nor its spatial organization has been previously described despite extensive anatomical and single-cell transcriptomic characterization of the species^33,71–73^.

Multiple independent observations support a close structural and functional association between flavinocytes and the central nervous system. Disruption of riboflavin uptake and metabolism impaired CNS regeneration (Fig. 7), whereas pharmacological and RNAi-mediated inhibition of neuronal signaling disrupted the re-establishment of the characteristic flavin-associated AF pattern, including the head-associated cluster and longitudinal strands (Figs. S7, S8). Moreover, flavinocyte-associated AF responded within seconds to chemical, physical, and light stimuli (Fig. 8A-C), while riboflavin pathway genes co-expressed with sensory markers within neural tissue (Fig. 8D), consistent with sensory-like characteristics.

Anatomically, flavinocytes form elongated longitudinal tracts immediately ventral to the cephalic ganglia and ventral nerve cords, originating from a prominent cluster in the head region (Fig. 2C,D). Their regularly spaced varicosities resemble the beaded morphology characteristic of neuronal processes, particularly those mediating volume transmission or neurosecretion rather than classical point-to-point synaptic communication^74^. Comparable varicose neurites have been extensively described across numerous invertebrate nervous systems, where regularly spaced varicosities serve as specialized sites of neurotransmitter and neuropeptide release^75–77^. Recent advances in tissue preservation and ultrastructural imaging have likewise revealed similar varicose neuronal morphologies in vertebrate nervous systems, including humans, suggesting that this architectural organization may be evolutionarily conserved^78,79^.

Assigning a positive identity, however, remains challenging: flavin-associated AF is lost upon chemical fixation^38^, whereas the planarian cell-type catalog rests on fixed material, so marker identity and flavin signal cannot be assessed in the same animal. Accordingly, classical panneuronal markers, including Synapsin and the planarian-specific axonal marker 1H6, do not label these structures. Given the limited availability of planarian-specific antibodies, we regard this negative result as inconclusive rather than evidence that flavinocytes are non-neuronal. Importantly, their processes contain tubulin in live planarians (Fig. S1), and directional transport is visible within the network (Movie 2), supporting the interpretation that flavinocytes represent an organized cellular structure rather than passive flavin accumulation.

Although excess riboflavin is cleared through excretion, several observations argue against an excretory identity. Neither fluorescent dextran labeling nor the protonephridial marker *Smed-inx10* overlapped with the flavinocyte network (Fig. S5). Moreover, the morphology of the two systems is fundamentally different: the protonephridial network forms a highly branched architecture terminating in dorsally positioned nephridiopores, whereas flavinocytes consist of largely unbranched longitudinal strands located ventrally^47,80^. In addition, flavin-associated AF was retained within the network after twelve weeks without food (Fig. 2B), which does not fit a protonephridial function in continuous ultrafiltration and solute clearance^81^. Comparable retention or storage of flavins has also been described in earthworms, where chloragocyte-derived eleocytes accumulate riboflavin within intracellular granules^82^, indicating that flavin-rich cells can serve a storage rather than a clearance function. Together, these observations establish flavinocytes as a cellular network distinct from the protonephridial system.

Interpretation of the potential cell identity is further complicated by the fact that cell type boundaries in early-diverging metazoans are less discrete than in vertebrates, with ancestral cells combining sensory, secretory, and metabolic functions that resolved only later in evolution into specialized cell types^83^. For example, nerveless sponges coordinate contractility through an ancient monoaminergic signaling system^84^. Likewise, concepts such as paraneurons, neurosecretory, and neuroendocrine cells, originally developed through studies of invertebrates and other early-diverging animals, highlight that sensory, secretory, metabolic, and neuronal characteristics can coexist within the same cell type rather than being partitioned into distinct cellular identities^85,86^. Consistent with this framework, the anatomical organization of flavinocytes, their close association with the CNS, stimulus-responsive behavior, and potential reciprocal functional relationship with the regenerating nervous system are compatible with a specialized neuro-associated cell population whose identity probably extends beyond conventional neuronal classifications.

Taken together, our findings support a model in which flavinocytes constitute a specialized neuro-associated network that spatially regulates vitamin B_2 a_vailability during regeneration. Their characteristic varicosities may function as specialized sites of localized flavin storage, transport, secretion, or metabolic exchange, thereby coordinating local cofactor availability with neuronal activity and the dynamic metabolic demands of neuronal remodeling during regeneration. Rather than relying solely on passive diffusion of water-soluble cofactors, regenerating tissues may actively organize flavin metabolism through dedicated cellular architectures that spatially match cofactor availability to local metabolic requirements. More broadly, our findings reveal that vitamin metabolism itself can be organized through specialized cellular networks, establishing local cofactor availability as an additional layer of metabolic regulation during tissue repair. Further unraveling the spatial organization of essential cofactors at the whole-organism level will provide fundamental insights into how metabolic architecture contributes to regeneration, and uncover conserved principles supporting tissue homeostasis, repair, and regeneration across animals.

## 4. Materials and methods

### 4.1 Planarian cultivation

An asexual strain of the freshwater planarian *Schmidtea mediterranea* was used for all experiments, unless otherwise specified. Animals were maintained in the dark at a constant temperature of 20°C in culture medium consisting of ultrapure H_2_O supplemented with 1.6 mM NaCl, 1 mM CaCl_2,_ 1 mM MgSO_4,_ 0.1 mM MgCl_2,_ 0.1 mM KCl, and 1.2 mM NaHCO_3._ Planarians were fed veal liver once per week and starved for at least 7 days before the start of each experiment. Animals receiving RF-related perturbations were starved for 3-4 weeks to promote a more riboflavin-depleted state. Regeneration was induced by amputation anterior to the pharynx, generating tail fragments that were used to study the effects of riboflavin-related perturbations on head regeneration, including the formation of a new brain. Specific experimental details are described in the corresponding methods section and figure legends.

### 4.2 Spectral autofluorescence characterization

To characterize endogenous autofluorescence (AF) and compare its spectral properties with those of synthetic flavins, spectral imaging was performed on a Zeiss LSM 880 confocal microscope (Carl Zeiss) in lambda mode at the Advanced Optical Microscopy Centre (UHasselt). Live planarians were immobilized with linalool as described in section 4.3. An excitation scan using the 458, 488, 514, 543, 594 and 633 nm laser lines was first performed to identify the excitation wavelength yielding the strongest AF signal, and 488 nm excitation was selected for spectral characterization. Lambda stacks were acquired with the 32-channel GaAsP detector from 414 to 690 nm in 8.9 nm channels, using an EC Plan-Neofluar 10×/0.30 objective for overview and tile-scan imaging of intact animals and an LD C-Apochromat 40×/1.1 W objective for cellular detail and synthetic compounds. Riboflavin (RF), flavin mononucleotide (FMN), flavin adenine dinucleotide (FAD), and nicotinamide adenine dinucleotide (NADH) were dissolved in planarian medium at 500 µM and imaged as droplets. Laser power, detector gain and pinhole were kept constant within each spectral comparison.

To visualize AF in the brain and ventral nerve cords, animals were exposed to 0.5 M MgCl₂ for 30 s prior to imaging. For each structure of interest (terminal cell bodies, varicosities, strands, and ganglia), at least three regions of interest (ROIs) per sample were drawn in ZEISS ZEN blue 3.7.7 (Carl Zeiss), and mean intensity per emission channel was exported. A background spectrum acquired outside the sample with identical settings was subtracted channel by channel, and spectra were normalized to their respective maximum. Representative spectra are shown; these were consistently observed across at least 8 control animals. Full spectral galleries are provided in Supplementary file 2.

### 4.3 Confocal and light-sheet imaging of live planarians

Live planarians were immobilized using the previously established linalool-based imaging approach^39^. Animals were exposed to 2.16 mM linalool prepared in fresh planarian medium and oriented according to the region of interest within a sealed FastWell chamber or FEP tube containing 2.16 mM linalool.

*In vivo* confocal imaging was performed using a Zeiss LSM 900 microscope (Carl Zeiss) equipped with an Axio Observer 7 platform, a Photometrics Prime 95B camera, an HXP 120 V light source, and GaAsP-PMT/multialkali PMT detectors. The optical setup used for live imaging included EC Plan-Neofluar 10×/0.30 and LD C-Apochromat 40×/1.1 W objectives. Fluorescence excitation was provided by 488 nm (10 mW), 561 nm (10 mW), and 641 nm (5 mW) diode lasers. Confocal images were acquired at 16-bit depth using a 1.0 AU pinhole.

Based on spectral characterization, endogenous flavin-associated AF was primarily visualized using 488 nm excitation. Whole-animal overview images used for quantitative AF measurements were acquired in widefield mode, whereas detailed visualization of flavinocyte-associated autofluorescent structures was performed in laser-scanning confocal (LSM) mode.

Live fluorescent labeling was performed for *in vivo* colocalization and anatomical characterization of flavin-associated AF. For visualization of microtubule organization, 6 µM Tubulin Tracker™ Deep Red (Thermo Fisher Scientific) was microinjected into the head mesenchyme (3× 32.2 nL) using a Nanoject II microinjector (Drummond Scientific), followed by incubation for 4 h in planarian medium prior to imaging. The excretory system was visualized by microinjection of 500-kDa fluorescent dextran (Invitrogen) into the head mesenchyme, as described previously^47^. To provide tissue-specific contrast of the intestine, animals were incubated with 0.5 µM dihydroethidium (DHE; Enzo Life Sciences) for 1 h in the dark, as described previously^87^. Following labeling, animals were washed twice in fresh planarian medium, immobilized, and imaged as described above.

For three-dimensional *in vivo* characterization and spatial localization of flavinocytes, volumetric light-sheet imaging was performed using a Zeiss Lightsheet 7 microscope (Carl Zeiss). Prior to imaging, live animals were incubated with Hoechst 33342 (Thermo Fisher Scientific; 1:5000) for 1 h to provide contrast for the body boundaries and subsequently immobilized using linalool. The microscope was equipped with two pco.edge 4.2 sCMOS cameras and an EC Plan-Neofluar 5×/0.16 detection objective. Samples were illuminated from both sides with a 7.98 µm light sheet (pivot scan on). Hoechst 33342 and endogenous flavin-associated AF were excited simultaneously with 405 nm and 488 nm lasers (50 mW each, at 10% power), and emission was split onto the two cameras using an LP 490 beam splitter. Z-stacks were acquired with a 3.76 µm step size as tiled scans with 10% overlap. Three-dimensional image processing, reconstruction, and visualization were performed using ZEISS arivis Pro 4.4. Identical acquisition settings were maintained between experimental groups within individual quantitative experiments.

### 4.4 Gene identification and single-cell transcriptomic data analysis

The publicly available single-cell RNA sequencing dataset from *Schmidtea mediterranea* generated by Fincher and colleagues was reanalyzed to investigate the cellular distribution of riboflavin uptake-and metabolism-related genes^44^. Gene identifiers were retrieved from PlanMine^88^ and S. Alvarado laboratory’s Rosetta Stone Transcript Mapper^89^ and are summarized in the Supplementary file 1, Table S1.

Expression of *Smed-RFT1*, *Smed-RFT2*, *Smed-RFK*, *Smed-FADS*, and *Smed-FMNr* was evaluated across anatomical regions, annotated cell types, and individual cell clusters. Pairwise co-expression analyses were performed between riboflavin pathway genes and selected marker genes representing neuronal subtypes, epidermal cells, neoblasts, cathepsin^+^/parenchymal cells, glial cells, protonephridial cells, and sensory cell populations. Heatmaps were generated using both average gene expression across all cells and average expression among expressing cells only (expression > 0). The number of expressing cells was annotated for each gene within each region, cell type, or cell cluster. Pairwise co-expression networks were generated to visualize the overlap between riboflavin pathway genes and lineage-specific marker genes. Data processing, visualization, and network analyses were performed in R (version 4.3.3)^90^, with specific libraries to preprocess the data matrices (tidyverse v2.0.0)^91^, to generate the heatmaps (ComplexHeatmap v2.18.0)^92^, and to draw the co-expression networks (ggraph v2.1.0; igraph v1.6.0)^93^. Force-directed Fruchterman and Reingold layout algorithm^94^ was used to distribute the nodes in the network graphs.

To validate these findings in independent datasets, the expression of riboflavin pathway genes was additionally examined in two other single-cell RNA sequencing atlases of *Schmidtea mediterranea.* First, scaled expression values for *Smed-RFT1*, *Smed-RFT2*, *Smed-RFK*, *Smed-FADS*, and *Smed-FMNr* across the 89 global cell clusters of the wild-type (WT) single-cell RNA sequencing dataset generated by Benham-Pyle and colleagues^46^ were obtained from the accompanying online data portal (https://simrcompbio.shinyapps.io/bbp_app/) and visualized as a heatmap, with clusters grouped by tissue according to the original cluster annotation. The heatmap was generated in R (version 4.4.1) using the pheatmap package (v1.0.13)^95^. Second, cell type-and cell subtype-specific expression profiles were obtained from the wild-type single-cell dataset (WT.sc.h5ad) of the 4D single-cell spatial transcriptomics atlas of Han and colleagues, using the PRISTA4D interactive portal^45^.

### 4.5 RNA interference

Gene knockdown was achieved by RNA interference (RNAi) using double-stranded RNA (dsRNA). dsRNA targeting the genes of interest was synthesized by *in vitro* transcription using the T7 RiboMAX™ Express RNAi System (Promega) according to the manufacturer’s instructions. Primer sequences are provided in Supplementary file 1, Table S2. Animals received six dsRNA injections over a two-week period (three consecutive daily injections per week for two weeks). Each injection consisted of 32.2 nL of 1.5 μg/μL dsRNA delivered ventrally between the photoreceptors and the pharynx using a Nanoject II microinjector (Drummond Scientific). Control animals were injected with an equal volume of ultrapure H₂O following the same injection schedule.

### 4.6 Whole-mount immunohistochemistry and fluorescent in situ hybridization

Whole-mount immunohistochemistry was performed to determine the localization of riboflavin transporter (RFT) relative to the planarian central nervous system using anti-SYNORF-1 as a pan-neural marker. Combined fluorescent *in situ* hybridization (FISH) and immunohistochemistry was additionally performed to examine the spatial relationship between RFT localization and neuronal populations expressing Smed-*chat*, *Smed-tph, Smed-vc-1*, and *Smed-pc2*.

For immunohistochemistry, the following primary antibodies were used: mouse anti-SYNORF-1 (Developmental Studies Hybridoma Bank; 1:10), and rabbit anti-RFT (Proteintech; 1:50). Alexa Fluor 488-and 568-conjugated secondary antibodies (Invitrogen; 1:500) were used for fluorescence detection. Whole-mount immunostaining was performed essentially as described previously^96^. Briefly, animals were euthanized and mucus was removed by immersion in 2% HCl for 5 min on ice. Samples were fixed in 4% formaldehyde prepared in PBSTx (1× PBS containing 0.3% Triton X-100) for 15 min at room temperature (RT), followed by overnight bleaching in 6% H₂O₂ in PBSTx under a cold white light source. Animals were blocked in 1% (w/v) bovine serum albumin (BSA) in PBSTx for 4 h at RT and incubated with the appropriate primary antibodies overnight (approximately 18 h) at 4°C. Following washing in PBSTx for 6–8 h, samples were incubated in blocking solution for 1 h at RT and subsequently with the appropriate Alexa Fluor-conjugated secondary antibodies for 16 to 18 h at 4°C.

For combined FISH and RFT immunostaining, FISH targeting the above-mentioned transcripts was performed first as described previously^97^. Gene-specific DIG-labeled RNA probes were generated using sequences of the gene fragments cloned in vectors, as summarized in Supplementary Table S3. Following the FISH protocol, samples were washed for 1 h in PBSTx and subjected to RFT immunostaining using the antibody and blocking incubation conditions described above. After immunostaining, samples were washed in PBSTx for at least 2 h and incubated with Hoechst 33342 (Invitrogen; 1:5,000) for 1 h at RT. Following a final wash in PBSTx, animals were mounted in ImmuMount (Thermo Fisher Scientific) and imaged using a Zeiss LSM 900 confocal microscope (Carl Zeiss).

### 4.7 Pharmacological interference

Pharmacological treatments were used to investigate whether neuronal signaling influences the spatial organization of flavin-associated AF during regeneration. Planarians were exposed to 500 μM tetrodotoxin (TTX; Enzo Life Sciences) to inhibit voltage-gated sodium channel-dependent neuronal activity; 117 μM 6-hydroxydopamine (6-OHDA; Merck) to disrupt catecholaminergic neurons; 1 mM 3-iodo-L-tyrosine (Thermo Fisher Scientific) to interfere with catecholamine synthesis; 10 μM fluoxetine (Merck) to modulate serotonergic signaling; 300 μM ondansetron (Merck) to antagonize 5-HT3 receptor-mediated serotonergic signaling; 300 μM atropine (Merck) to inhibit muscarinic cholinergic signaling; and 300 μM hexamethonium (Merck) to inhibit nicotinic cholinergic signaling. 0.025 μM Rotenone (Sigma-Aldrich) was used to interfere with mitochondrial electron transport through inhibition of complex I.

Compounds were prepared in planarian cultivation medium. Animals were pre-exposed for 1 day prior to amputation and subsequently exposed throughout 7 days of regeneration, with treatment solutions renewed every other day. Following treatment, animals were washed in fresh planarian medium, and flavin-associated AF was assessed by *in vivo* imaging as described above.

### 4.8 qPCR

Knockdown efficiency of *Smed-RFT1*, *Smed-RFT2*, *Smed-RFK*, *Smed-FADS*, and *Smed-FMNr* was assessed in intact animals following RNAi treatment and prior to amputation. Four biological replicates were collected per experimental condition. Samples were snap-frozen and lysed in 100 μL of Qiagen lysis buffer containing 1% β-mercaptoethanol. RNA was extracted using a standard phenol–chloroform protocol and precipitated using sodium acetate and ethanol. RNA concentration and purity were assessed using a NanoDrop ND-1000 spectrophotometer, after which genomic DNA was removed using the TURBO DNA-free™ Kit (Invitrogen). cDNA was synthesized using the SuperScript™ III First-Strand Synthesis SuperMix (Thermo Fisher Scientific), diluted 1:9, and analyzed by quantitative real-time PCR using a QuantStudio™ 5 Real-Time PCR System (Thermo Fisher Scientific). *Smed-GUSB* and *Smed-HMBS* were selected as stable reference genes using the geNorm algorithm in qbase+ (Biogazelle) and used for normalization. Primer sequences are provided in Table S4.

### 4.9 Characterization of planarian behavior and tissue regeneration phenotypes

Locomotor activity was assessed using a grid-based movement assay adapted from Hagstrom and colleagues^98^. Individual planarians were transferred to Petri dishes containing cultivation medium and positioned above a 0.5-cm grid. Animals were allowed to acclimatize for 20 s, after which movement was monitored for 60 s. Locomotor activity was determined from the total number of grid lines crossed during this period.

Regenerative growth was evaluated at 7 days post-amputation (7 DPA) by quantifying newly formed blastema tissue. Live regenerating fragments were imaged using a Nikon SMZ800 stereomicroscope equipped with a Nikon DS-Ri2 digital camera. Blastema and whole-animal areas were delineated in Fiji (ImageJ)^99^, and relative blastema size was calculated as the blastema area divided by the total body area.

Eye regeneration was evaluated at 7 DPA by visual inspection of live animals under bright-field illumination at high magnification and across multiple focal planes. Regenerating animals were assigned to one of four phenotypic categories: two normally developed eyes, two faint or incompletely developed eyes, one eye, or complete absence of visible eyes.

Central nervous system regeneration was assessed at 7 DPA following whole-mount anti-SYNORF-1 immunostaining as described above. Brain development was quantified using two complementary morphological parameters in Fiji (ImageJ). The total area occupied by the bilobed cephalic ganglia was delineated and normalized to the total planarian body area. Anterior commissure regeneration was additionally evaluated by scoring animals according to the presence or absence of a continuous synapsin-positive connection between the two cephalic ganglia.

For all representative phenotype images, *n* represents the total number of animals showing the phenotype per experimental condition. Images were selected to illustrate the predominant phenotype observed within each group, while the accompanying quantitative analyses provide further insight into phenotypic variability.

### 4.10 Exposure to different stress stimuli

The response of flavin-associated AF to chemical and physical stimuli was assessed in vivo. Planarians were exposed to 0.2% chlorotone for 10 min, or 2.5% polystyrene particles (0.1 mm diameter) during imaging. Following exposure, animals were imaged as described above to assess phenotypical changes in the spatial organization of flavin-associated AF. To investigate the response to light stimulation, live planarians were continuously exposed to 488 nm excitation for 1 h while AF was recorded by time-lapse imaging at 10-s intervals.

### 4.11 Statistics

All statistical analyses were performed using RStudio (version 2024.04.2+764). Data normality was assessed using the Shapiro–Wilk test. For comparisons between independent groups, an unpaired Student’s t-test was used when the assumption of normality was met. When normality was not met, including after square-root or log transformation where appropriate, the non-parametric Wilcoxon rank-sum test was applied. All comparisons were performed against the respective control groups, with Holm correction applied to account for multiple testing where applicable. *P* values < 0.05 were considered statistically significant. Quantitative data are presented as the mean ± SEM; categorical phenotypes are presented as percentages.

## Acknowledgments

We are grateful to Lukas Schärer for the valuable scientific discussions. M.H. thanks Sam Duwe for his guidance and training in microscopy techniques. We thank Natascha Steffanie and Ria Vanderspikken for their skillful technical assistance. Renée Moonen, Lucia Malikova, Wout Vanbuel, Dillan Winters, and Thijmen Isken are thanked for their contributions during their internships.

## Author contributions

**M.H**.: conceptualization, data curation, formal analysis, investigation, methodology, supervision, visualization, writing-original draft, writing-review and editing. **V.J.:** investigation, methodology. **R.M.:** data curation, investigation, methodology. **J.F.A.:** formal analysis, visualization, writing-review and editing. **R.V.:** data curation, investigation, writing-review and editing. **C.S.:** investigation, writing-review and editing. **T.A:** conceptualization, writing-review and editing. **N.D.S.:** methodology, writing-review and editing. **F.C.:** conceptualization, resources, writing-review and editing. **K.S:** conceptualization, funding acquisition, project administration, resources, supervision, writing-review and editing.

## Data availability statement

All single-cell data analyzed in this study are publicly available: single-cell RNA sequencing data from Fincher et al.^44^, Benham-Pyle et al.^46^ (https://simrcompbio.shinyapps.io/bbp_app/), and Han et al. (PRISTA4D)^45^. R code used for processing and visualization of the Fincher et al. dataset has been deposited in Zenodo (https://doi.org/10.5281/zenodo.23266279). All other relevant data are included in the manuscript and supplementary files.

## Supplemental information

Supplementary file 1: Figures S1-S9 and Tables S1-S4.

Supplementary file 2: Representative Spectral Galleries; Figures S10-S14.

## Supplemental movies

Movie 1. Three-dimensional lightsheet reconstruction of the flavinocyte network in a live intact planarian *Schmidtea mediterranea.* Endogenous flavin-associated autofluorescence is shown in yellow/orange, while Hoechst 33342 fluorescence was used to generate a 3D surface rendering of the animal (blue), illustrating the spatial distribution of the flavinocyte network within the intact organism (scale, 50 μm).

Movie 2. Dynamic movement of flavin-associated autofluorescence along longitudinal flavinocyte strands in live *Schmidtea mediterranea.* Time-lapse wide-field imaging showing dynamic transport of autofluorescent cargo along the longitudinal flavinocyte network. (40x water-immersion objective; scale, 50 μm)

