## Supplementary file 2 for "Flavins in Focus: Vitamin B_2_ as a Metabolic Keystone for Neuronal Repair in Regenerating Planarians"

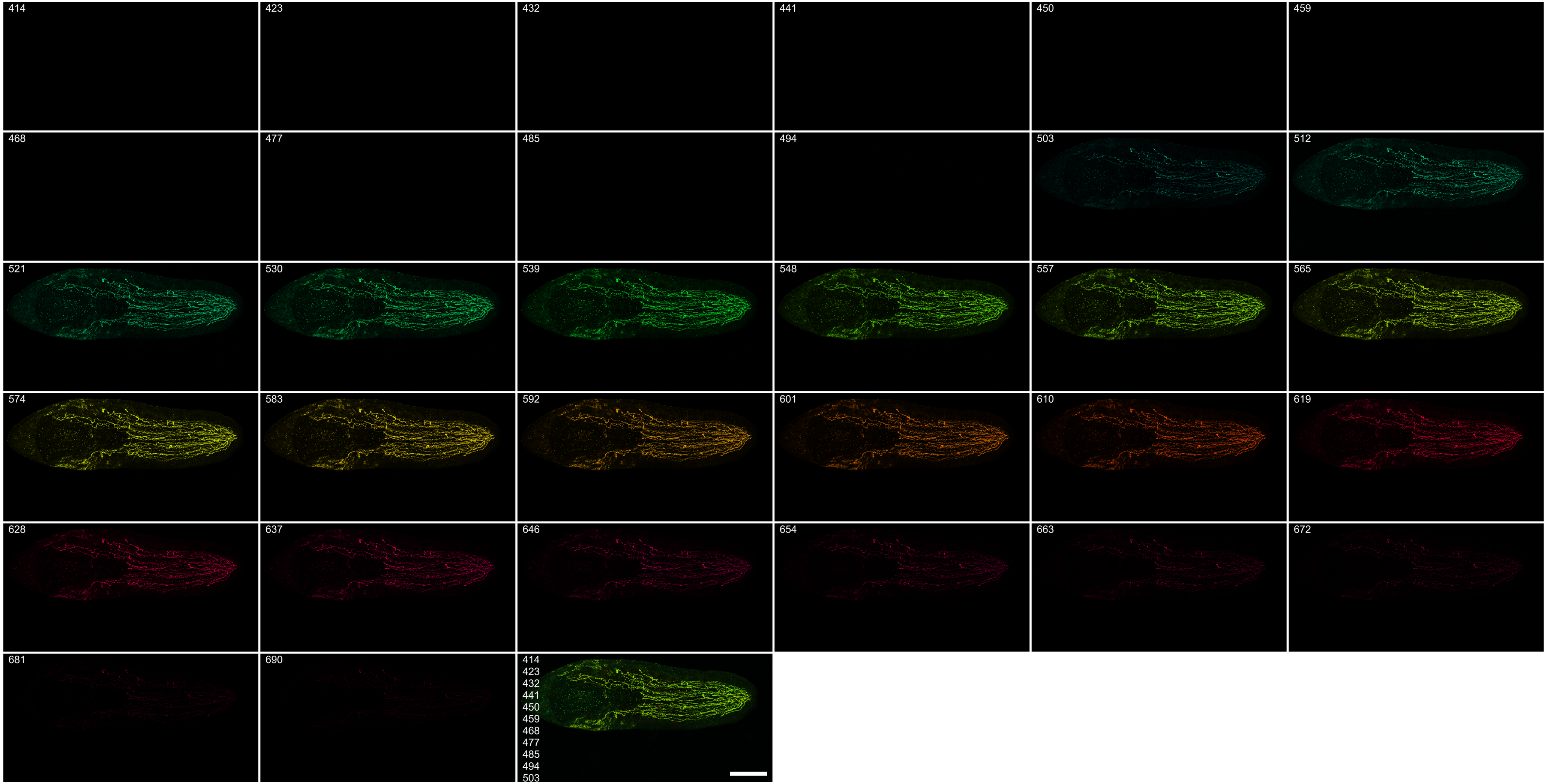


**Figure S10. Spectral gallery of an intact *Schmidtea mediterranea planarian in vivo.*** Lambda stack acquired with 488 nm excitation (10× objective); emission was detected in 32 channels of ~9 nm spanning 414-690 nm. The final panel shows the overlay of all channels. Display settings are identical across all panels. Scale bar, 500 µm (applies to all panels).

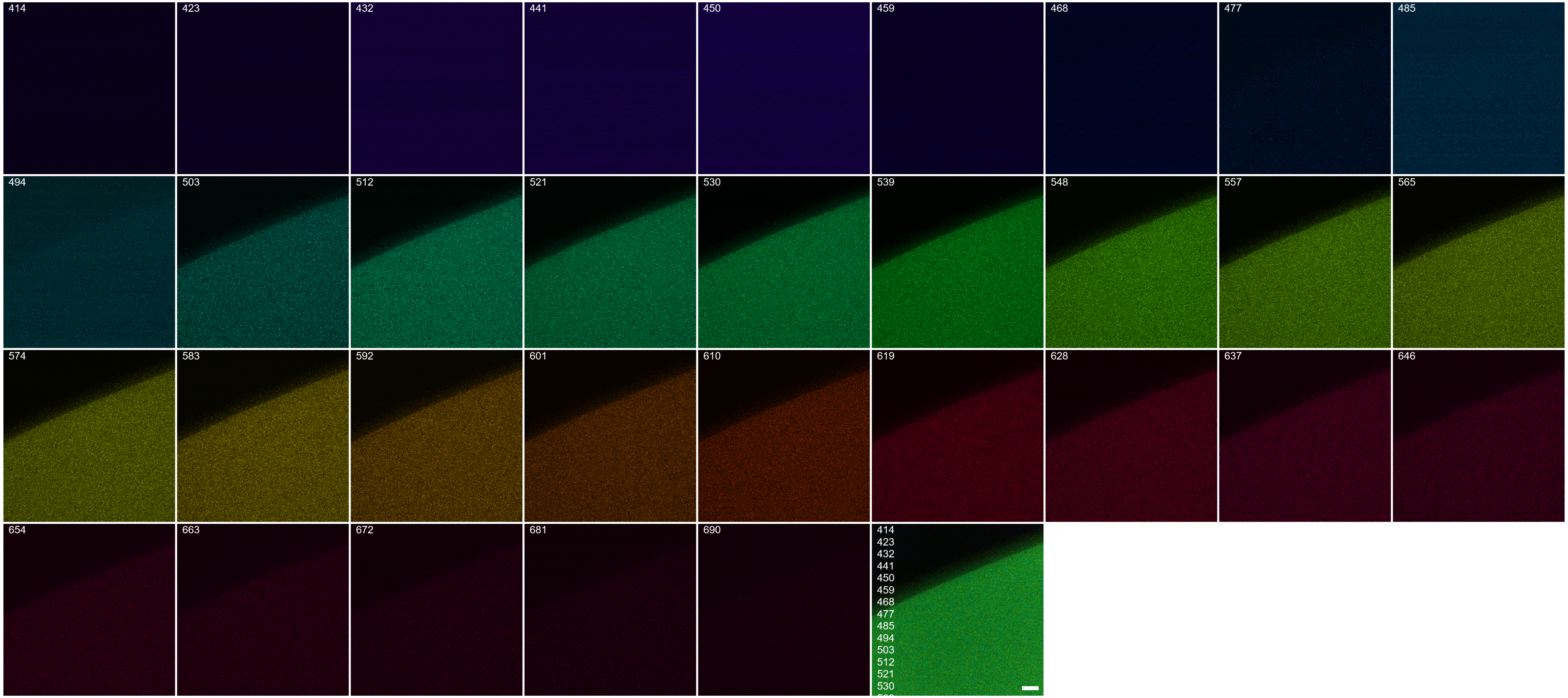

**Figure S11. Spectral gallery of a 500 µM synthetic riboflavin droplet.** Lambda stack acquired with 488 nm excitation (40× water-immersion objective); emission was detected in 32 channels of ~9 nm spanning 414-690 nm. The final panel shows the overlay of all channels. Display settings are identical across all panels. Scale bar, 20 µm (applies to all panels).

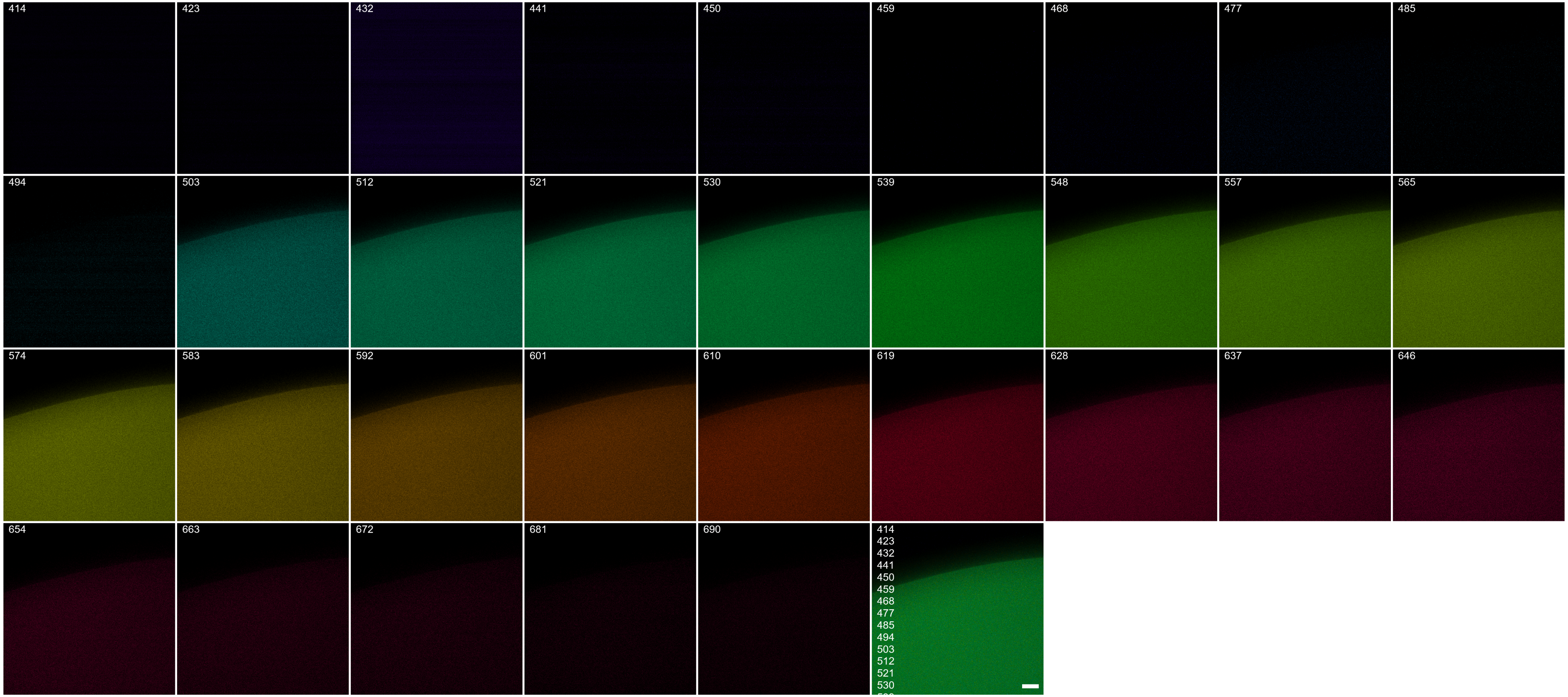


**Figure S12. Spectral gallery of a 500 µM synthetic FMN droplet.** Lambda stack acquired with 488 nm excitation (40× water-immersion objective); emission was detected in 32 channels of ~9 nm spanning 414-690 nm. The final panel shows the overlay of all channels. Display settings are identical across all panels. Scale bar, 20 µm (applies to all panels).

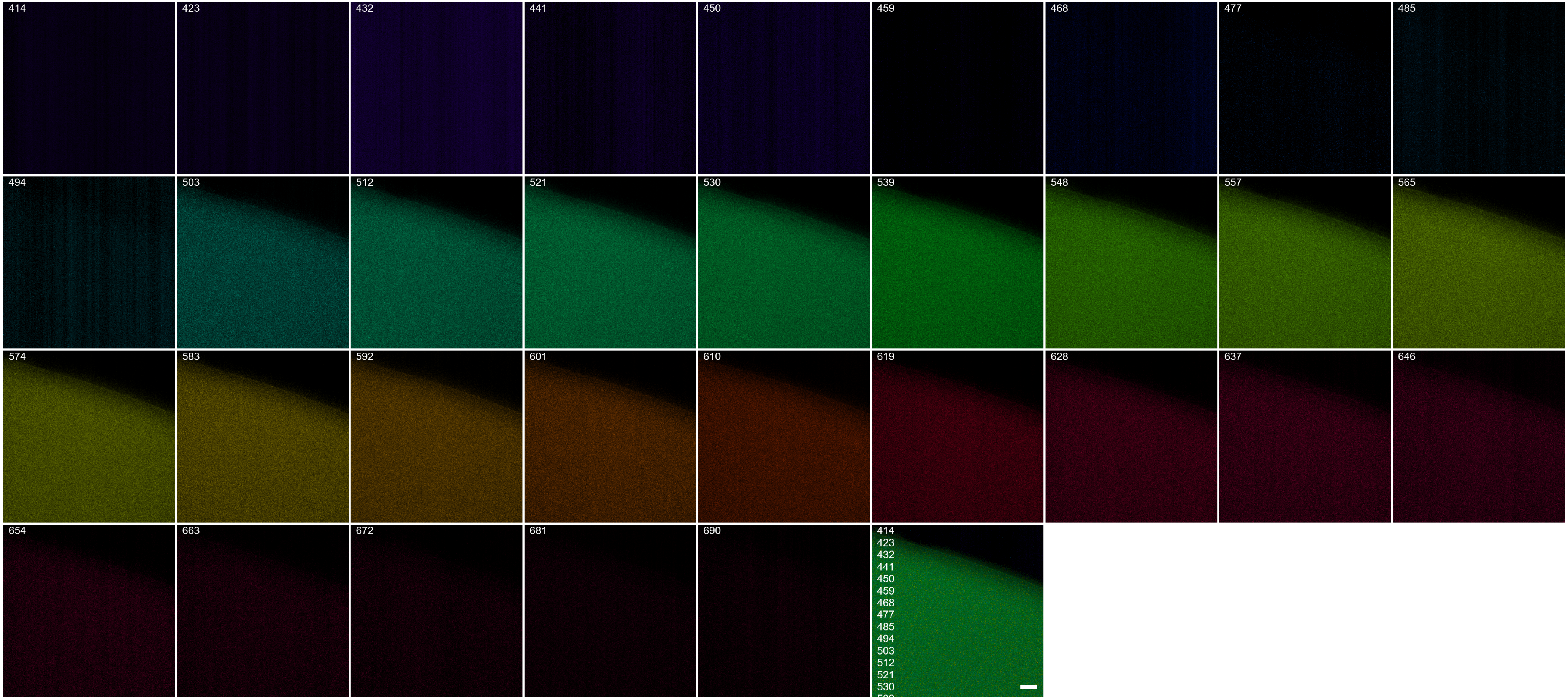


**Figure S13. Spectral gallery of a 500 μM synthetic FAD droplet.** Lambda stack acquired with 488 nm excitation (40× water-immersion objective); emission was detected in 32 channels of ~9 nm spanning 414-690 nm. The final panel shows the overlay of all channels. Display settings are identical across all panels. Scale bar, 20 μm (applies to all panels).

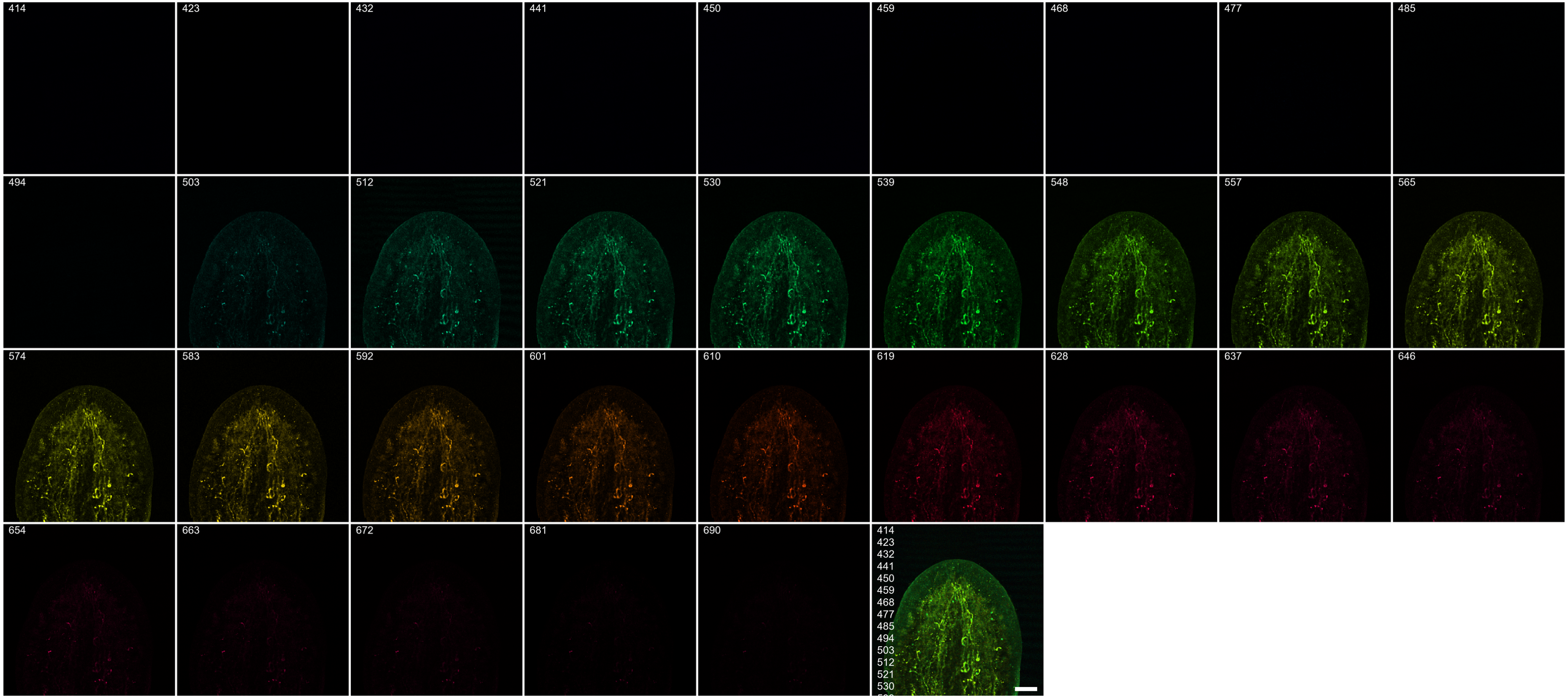


**Figure S14. Spectral gallery of the head region of an intact *Schmidtea mediterranea* planarian *in vivo* following exposure to 0.5 M $MgCl_2$ for 30 s.** Lambda stack acquired with 488 nm excitation (10× objective); emission was detected in 32 channels of ~9 nm spanning 414-690 nm. The final panel shows the overlay of all channels. Display settings are identical across all panels. Scale bar, 200 μm (applies to all panels).
