## Supplemental file 1 for "Flavins in Focus: Vitamin B_2_ as a Metabolic Keystone for Neuronal Repair in Regenerating Planarians"

### Supplemental Figures

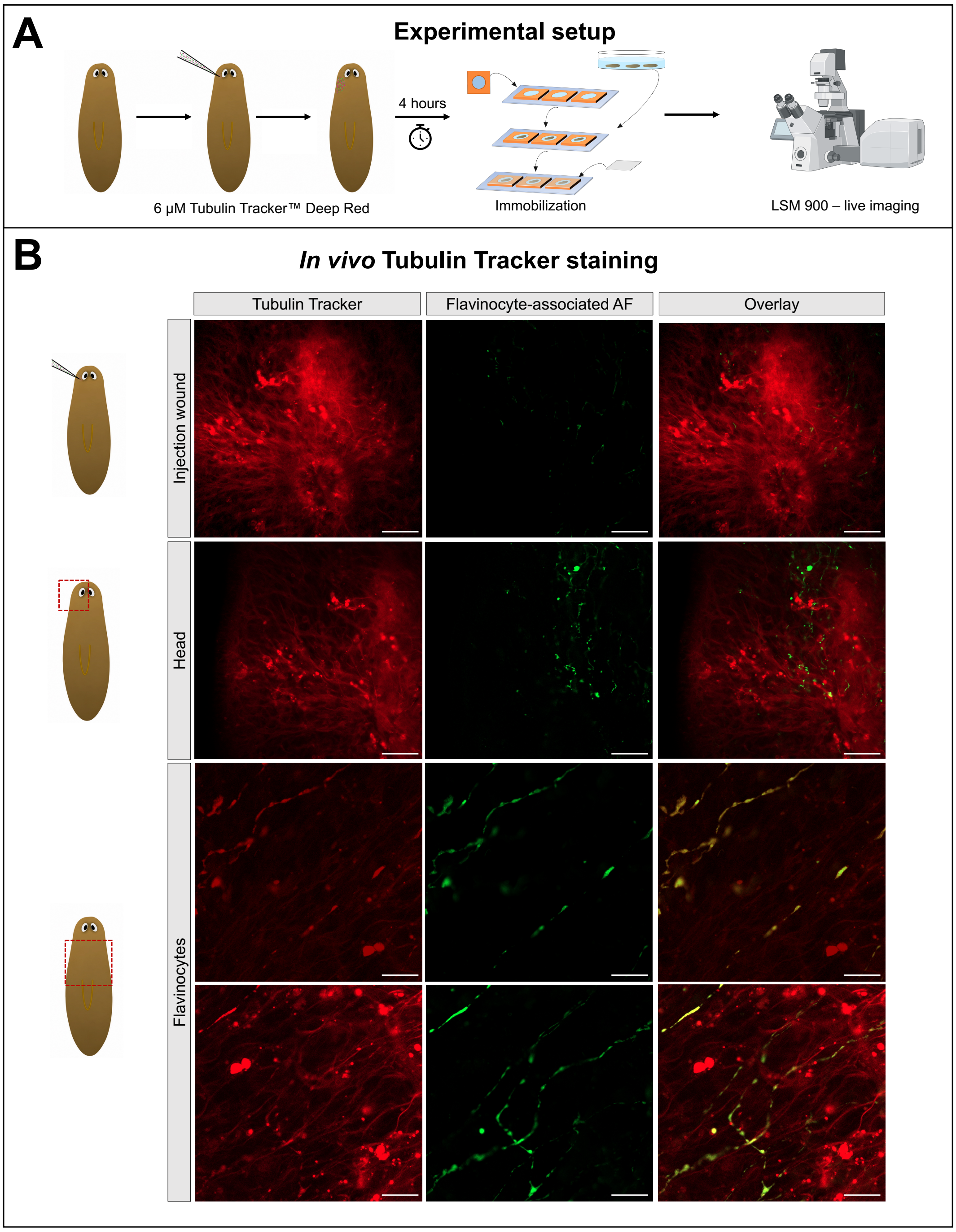


**Figure S1. Flavinocyte-associated autofluorescence colocalizes with tubulin-containing processes *in vivo*. A)** Graphical representation of the experimental setup. Live planarians were microinjected with 6 µM Tubulin Tracker™ Deep Red into the head mesenchyme, incubated for 4 h in medium, immobilized using linalool, and imaged using a Zeiss LSM900 confocal microscope. Created in BioRender. Heleven, M. (2026) https://BioRender.com/1dxl2ep. **B)** Representative live images showing Tubulin Tracker labeling (red), flavinocyte-associated autofluorescence (green), and merged images in the injection region, head, and longitudinal flavinocytes. Tubulin Tracker labeling overlapped with flavinocyte-associated AF along elongated strands and varicosity-like structures (scale bars, 10 µm). Images are representative of three independent biological replicates.

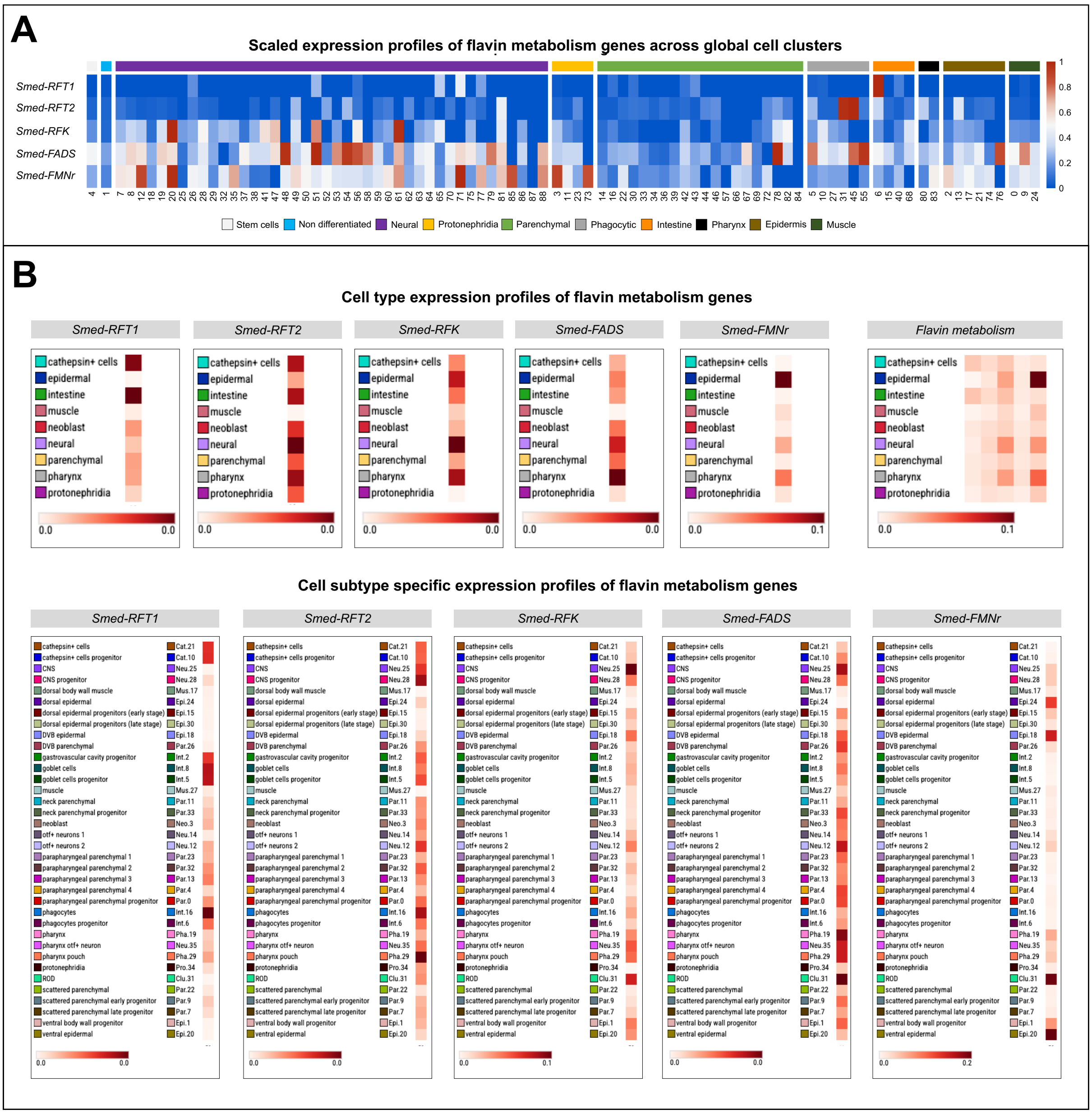


**Figure S2. Riboflavin-related gene expression across independent single-cell transcriptomic datasets. A)** Scaled expression profiles of *Smed-RFT1*, *Smed-RFT2*, *Smed-RFK*, *Smed-FADS*, and *Smed-FMNr* across global cell clusters from the single-cell RNA-sequencing dataset of Benham-Pyle et al. (2021). Cell clusters are grouped into major cell classes, including stem cells, non-differentiated cells, neural cells, protonephridia, parenchymal cells, phagocytic cells, intestine, pharynx, epidermis, and muscle. **B)** Cell type and cell subtype expression profiles of *Smed-RFT1*, *Smed-RFT2*, *Smed-RFK*, *Smed-FADS*, and *Smed-FMNr* from the spatial single-cell transcriptomic dataset of Han et al. (2026). Upper panels show expression across major cell types and the combined expression profile of the five riboflavin-related genes ("Flavin metabolism"), while lower panels show gene-specific expression across transcriptionally defined cell subtypes.

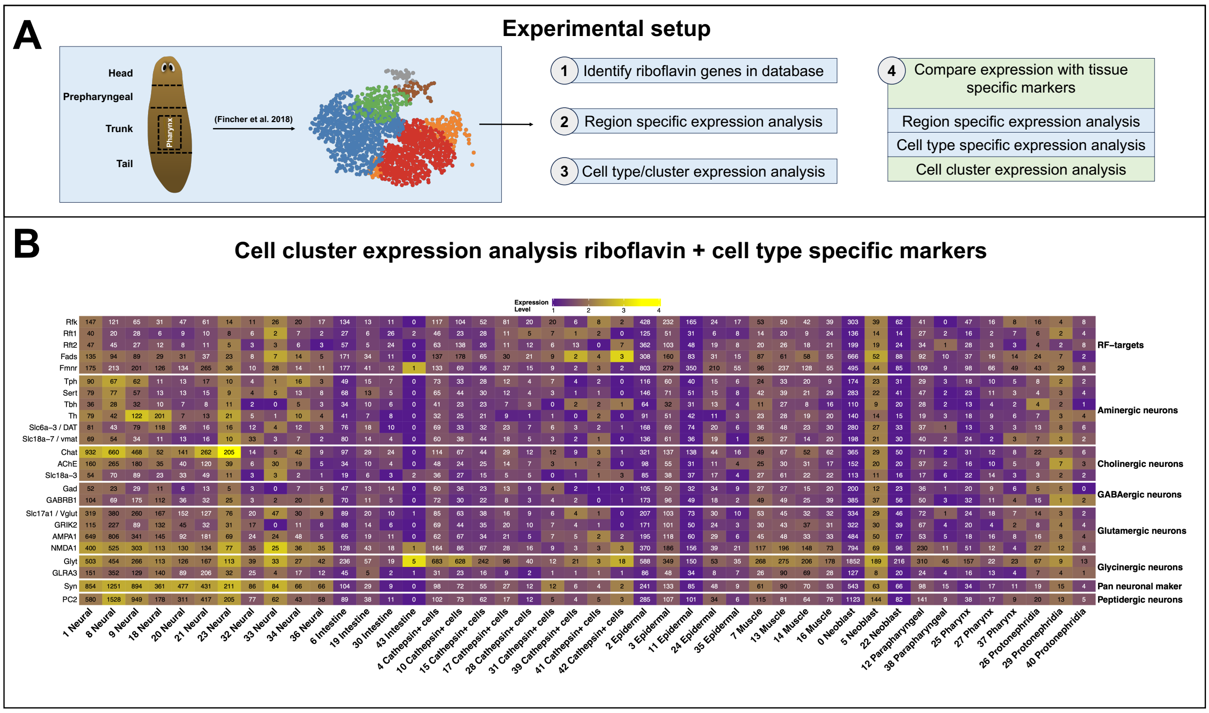


**Figure S3. Comparative cell cluster analysis of RF machinery and neuronal markers. A)** Graphical representation of the experimental setup. The published single-cell RNA-sequencing data from Fincher et al. (2018) were analyzed to characterize the expression of genes involved in riboflavin uptake and metabolism within individual cell clusters. Created in BioRender. Heleven, M. (2026) https://BioRender.com/1dxl2ep. **B)** Cell cluster-specific expression of *Smed-RFT1*, *Smed-RFT2*, *Smed-RFK*, *Smed-FADS*, and *Smed-FMNr* compared with markers of aminergic, cholinergic, GABAergic, glutamatergic, glycinergic, and peptidergic neurons.

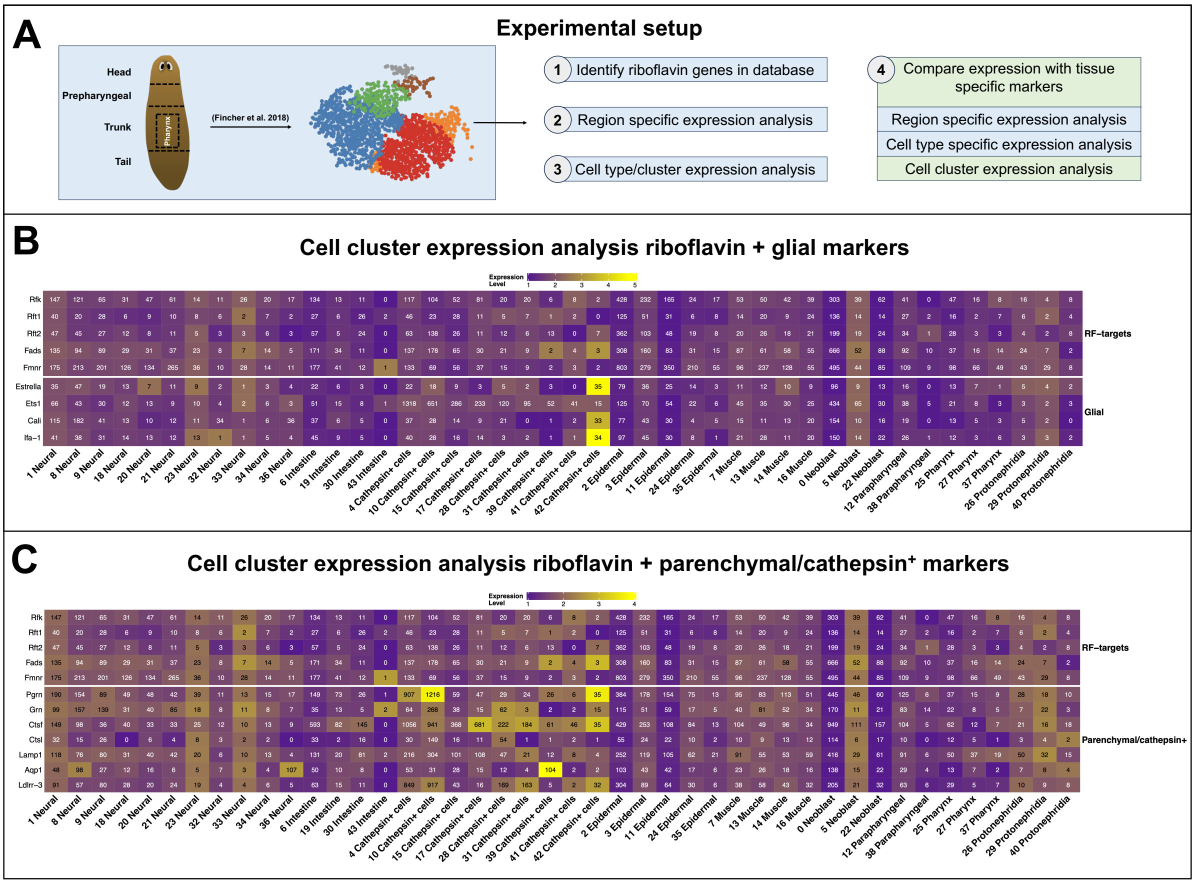


**Figure S4. Comparative cell cluster analysis of riboflavin machinery and glial/parenchymal markers. A)** Graphical representation of the experimental setup. Published single-cell RNA-sequencing data from Fincher et al. (2018) were analyzed to characterize the expression of genes involved in riboflavin uptake and metabolism within individual cell clusters. Created in BioRender. Heleven, M. (2026) https://BioRender.com/1dxl2ep. **B)** Cell cluster-specific expression of *Smed-RFT1*, *Smed-RFT2*, *Smed-RFK*, *Smed-FADS*, and *Smed-FMNr* compared with glial markers. **C)** Cell cluster-specific expression of *Smed-RFT1*, *Smed-RFT2*, *Smed-RFK*, *Smed-FADS*, and *Smed-FMNr* compared to parenchymal/cathepsin$^{+}$ markers.

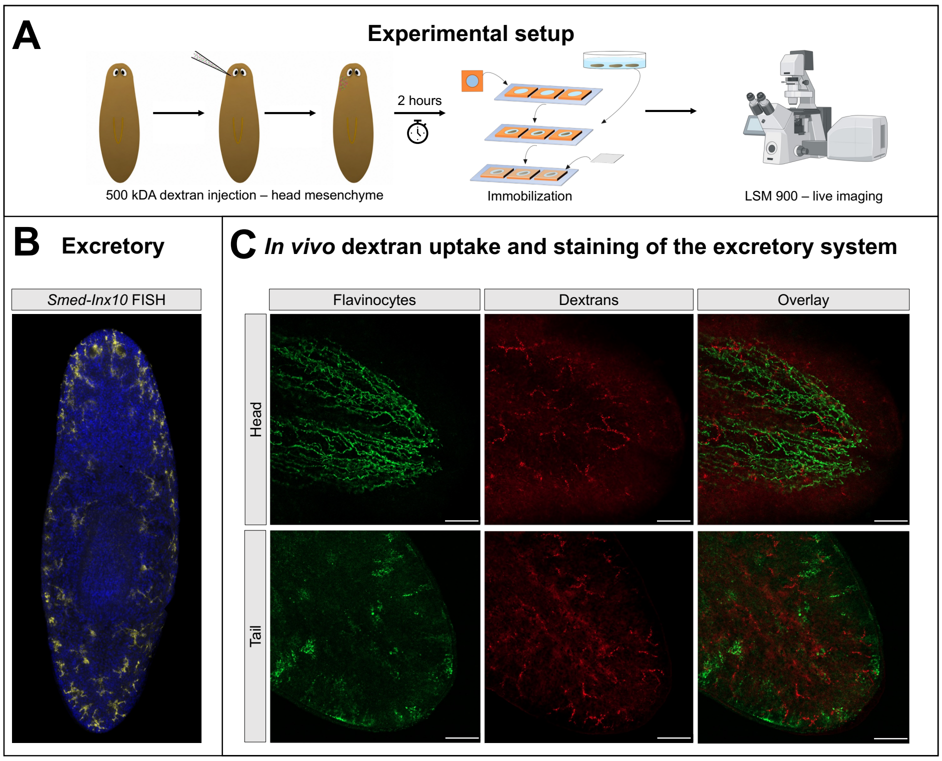


**Figure S5. Flavinocyte-associated autofluorescence is spatially distinct from dextran-labeled excretory structures. A)** Graphical representation of the experimental setup. Animals received three head mesenchymal injections of 32.2 nL of 500 kDa dextran, were incubated for 2 h, immobilized, and imaged live using a Zeiss LSM900 confocal microscope. Created in BioRender. Heleven, M. (2026) https://BioRender.com/1dxl2ep. **B)** Whole-mount fluorescence in situ hybridization (FISH) of the excretory marker *Smed-inx10* in intact animals. **C)** Representative live images of endogenous flavin autofluorescence (green) and 500-kDa dextran labeling of the excretory system (red). Single-channel images and corresponding merged images are shown for the head and tail regions (scale bars, 100 µm).

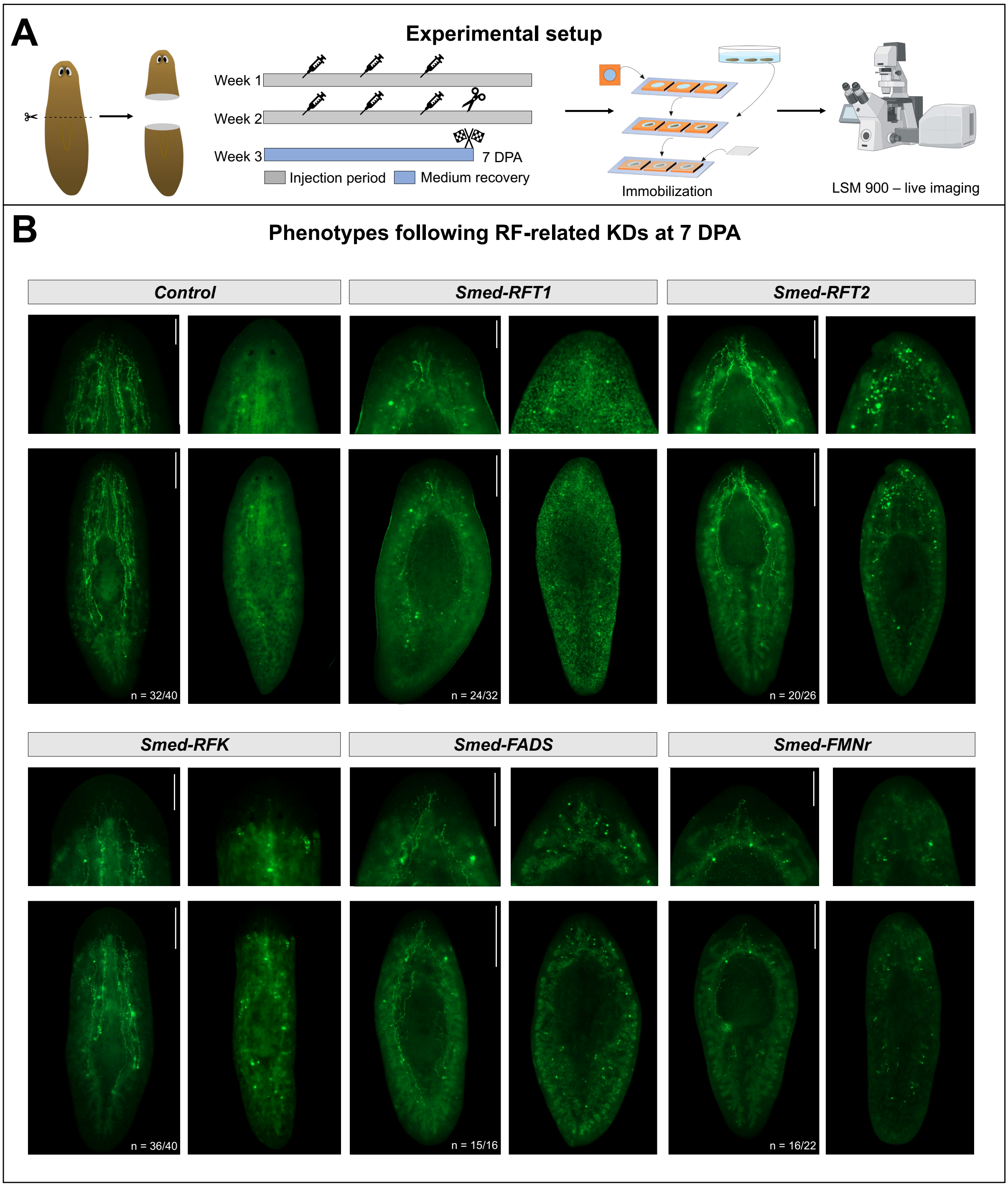


**Figure S6. Riboflavin-related gene knockdown alters flavin autofluorescence in regenerating tails. A)** Graphical representation of the experimental setup. Animals were subjected to two rounds of RNA interference (RNAi), each consisting of three injection days, targeting riboflavin-related genes. Following RNAi, animals were amputated prior to the pharynx and allowed to regenerate. At 7 days post-amputation (7 DPA), planarians were immobilized for live imaging using a Zeiss LSM900 confocal microscope. Created in BioRender. Heleven, M. (2026) https://BioRender.com/1dxl2ep. **B)** Representative live images following RNAi of *Smed-RFT1*, *Smed-RFT2*, *Smed-RFK*, *Smed-FADS*, and *Smed-FMNr* are shown. Whole-animal views (scale bars, 500 µm) and zoomed-in insets (scale bars, 250 µm) of the head region are shown from both dorsal and ventral sides. Autofluorescence (AF) display intensity was adjusted independently between images to visualize differences in AF distribution; images should therefore not be used for direct intensity comparison. The indicated *n* represents the total number of animals showing the respective phenotype across more than three independent experiments, except for *Smed-FADS* RNAi, which was assessed across two independent experiments.

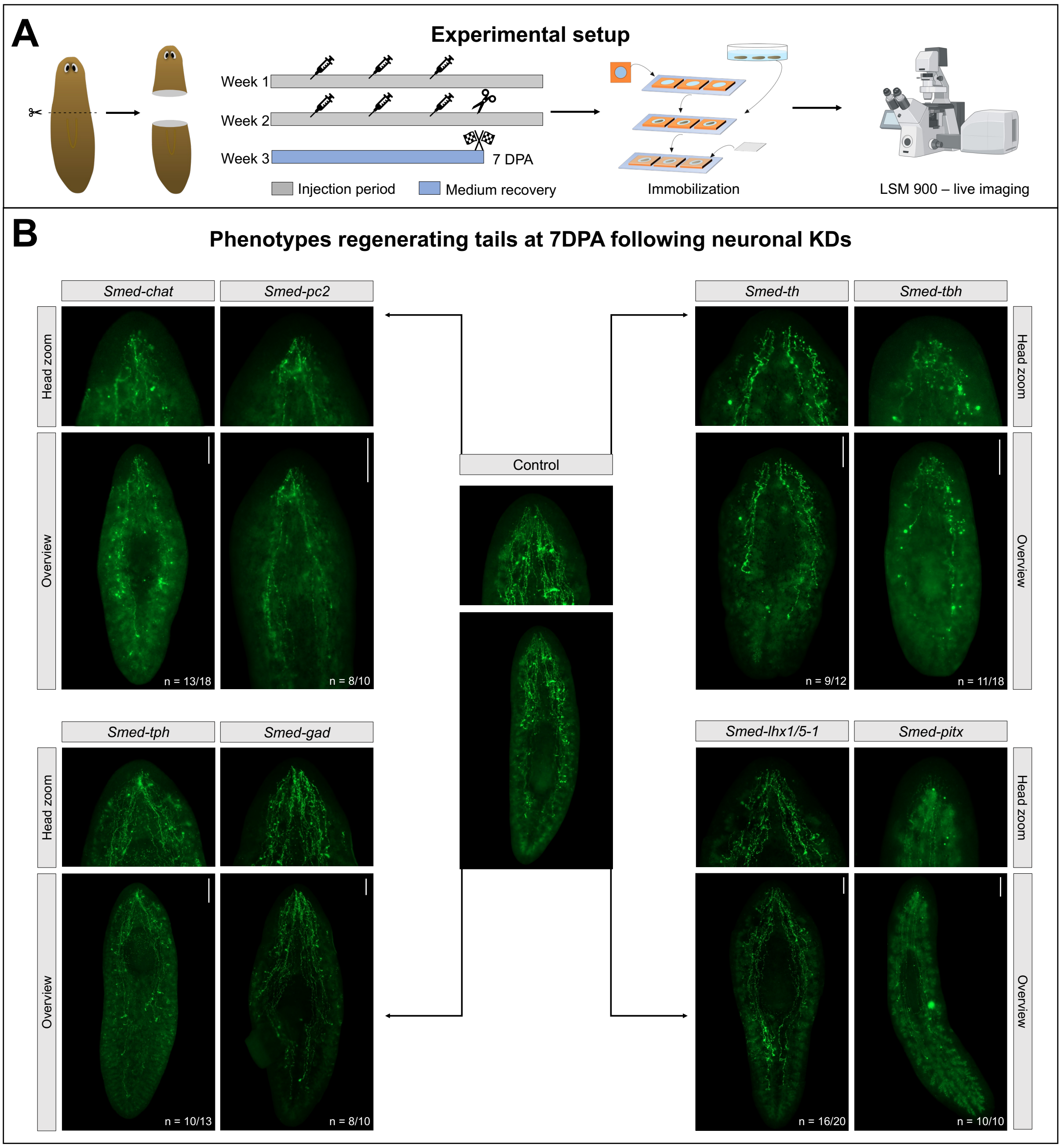


**Figure S7. Neuronal marker knockdown affects the spatial distribution of flavin-associated AF. A)** Graphical representation of the experimental setup. Animals were subjected to two rounds of RNA interference (RNAi), each consisting of three injection days, targeting neuronal markers. Following RNAi, animals were amputated prior to the pharynx and allowed to regenerate. At 7 days post-amputation (7 DPA), planarians were immobilized for live imaging using a Zeiss LSM900 confocal microscope. Created in BioRender. Heleven, M. (2026) https://BioRender.com/1dxl2ep. **B)** Representative live images of regenerating tails following control RNAi or RNAi of *Smed-chat*, *Smed-pc2*, *Smed-th*, *Smed-tbh*, *Smed-tph*, *Smed-gad*, *Smed-lhx1/5-1*, and *Smed-pitx* at 7 DPA. Whole-animal overviews and zoomed-in insets of the regenerated head region are shown (scale bars, 200 µm). The indicated *n* represents the total number of animals showing the respective phenotype across two independent experiments, except for *Smed-pc2 and Smed-pitx* RNAi, which were assessed in one experiment.

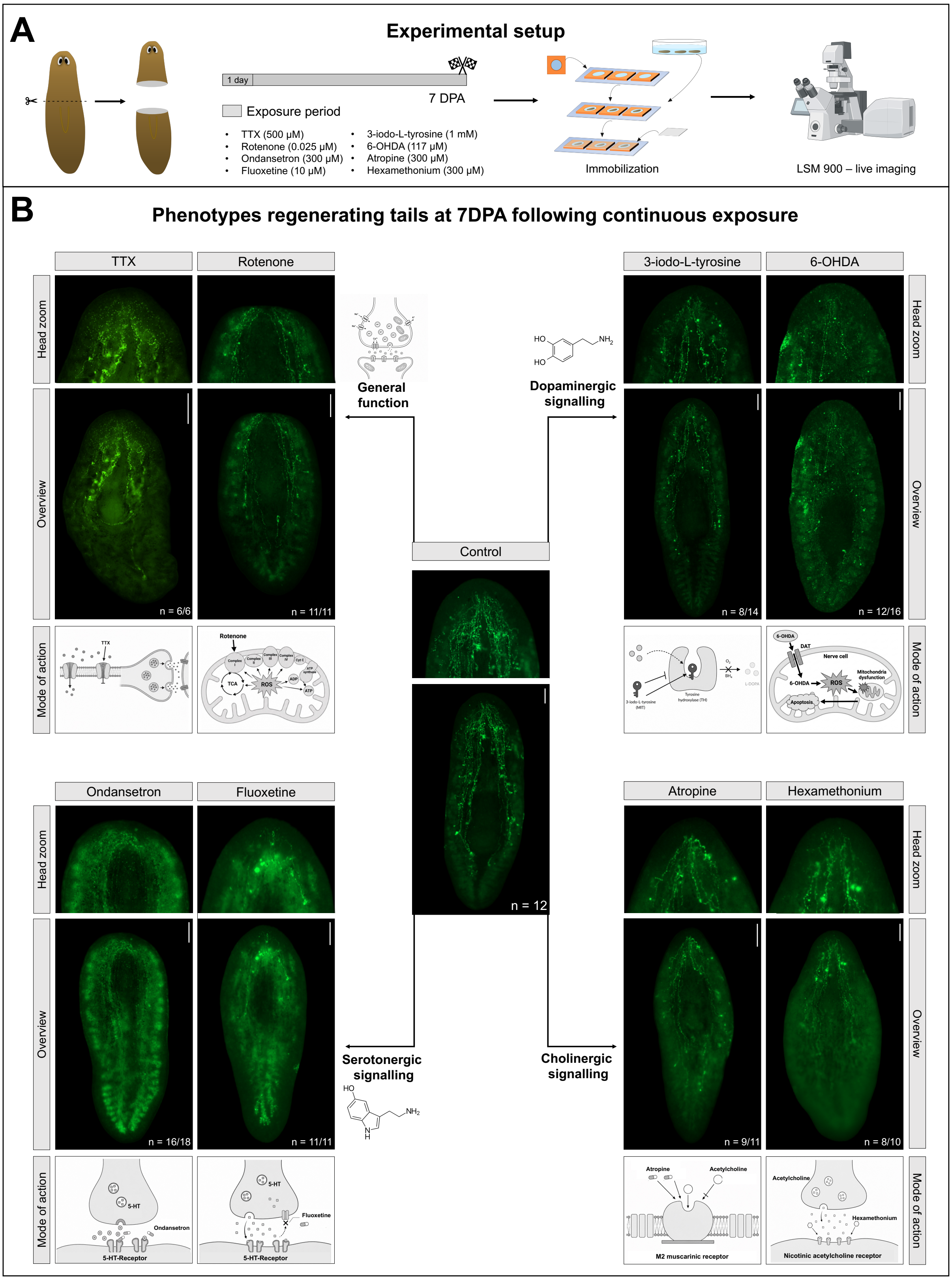


**Figure S8. Pharmacological interference impairs the spatial distribution of flavin-associated AF. A)** Graphical representation of the experimental setup. Animals were pre-exposed to pharmacological compounds for 1 day before being amputated prior to the pharynx. Animals remained continuously exposed throughout regeneration, and live imaging was performed at 7 days post-amputation (7 DPA) using a Zeiss LSM900 confocal microscope. Created in BioRender. Heleven, M. (2026) https://BioRender.com/1dxl2ep. **B)** Representative live images of regenerating tails following continuous exposure to compounds targeting general neuronal function (tetrodotoxin and rotenone), dopaminergic signaling (3-iodo-L-tyrosine and 6-hydroxydopamine), serotonergic signaling (ondansetron and fluoxetine), and cholinergic signaling (atropine and hexamethonium) are shown. Representative control animals are shown in the center of the panel. Whole-animal overviews and zoomed-in insets of the regenerated head region are shown (scale bars, 200 μm), alongside schematic illustrations of the corresponding mechanisms of action. The indicated *n* represents the total number of animals showing the respective phenotype across two independent experiments, except for hexamethonium, fluoxetine, and atropine, which were assessed in one experiment.

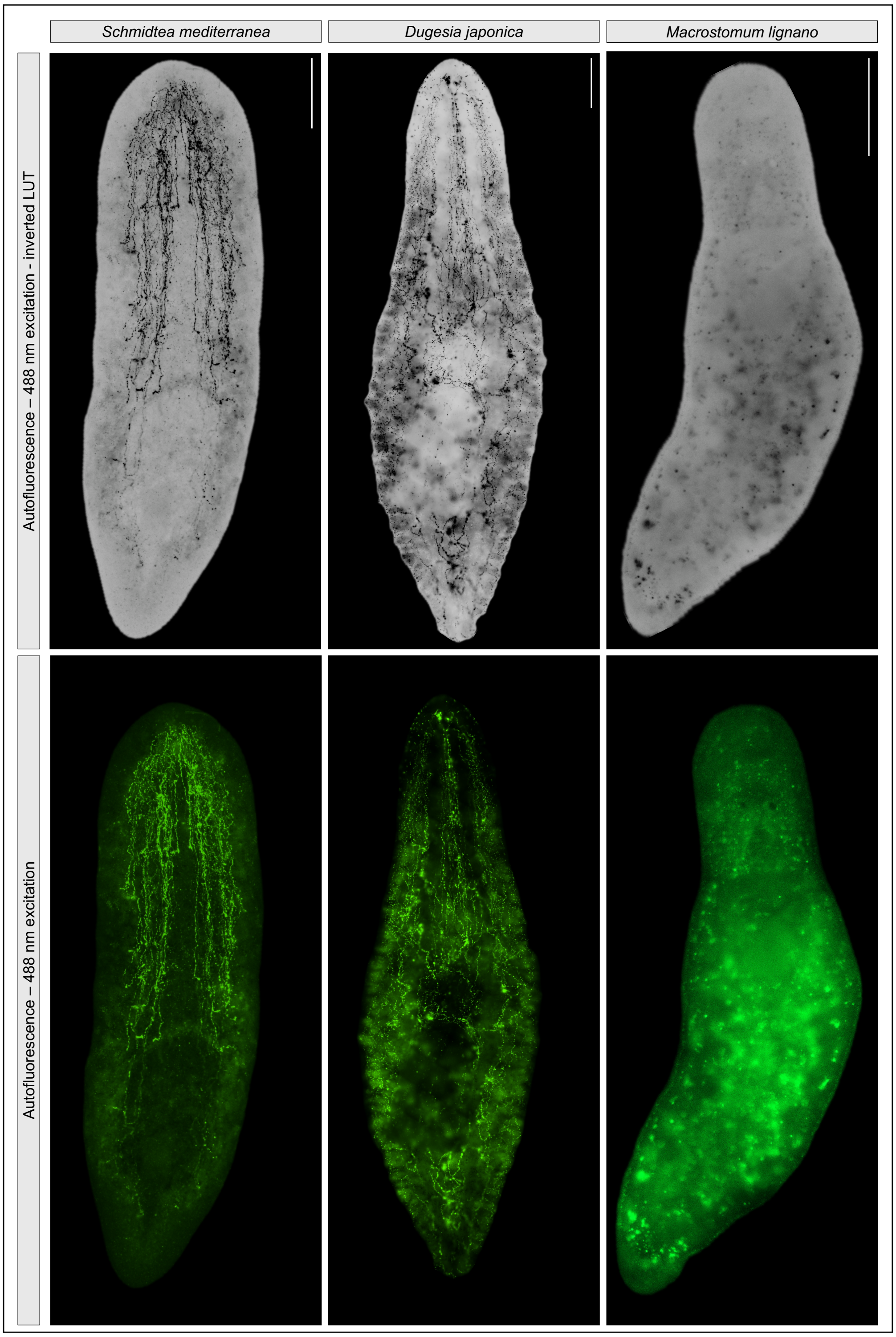


**Figure S9. Comparative analysis of flavinocyte-associated autofluorescence across flatworm species.** Representative live images of endogenous autofluorescence in the asexual planarian *Schmidtea mediterranea* (scale bar, 500 µm), asexual planarian *Dugesia japonica* (scale bar, 500 µm), and the flatworm *Macrostomum lignano* (scale bar, 100 µm), acquired using 488 nm excitation. Images are presented using inverted LUT (upper panels) and green autofluorescence (lower panels).

### Supplemental Tables

**Table S1: Gene identifiers of riboflavin-related genes in *Schmidtea mediterranea***

| Gene | PlanMine | Benham-pyle et al. 2021 | Han et al. 2026 |
|---|---|---|---|
| *Smed-RFT1* | dd_Smed_v6_11730_0_1 | SMED30034985 | SMESG000040849.1 |
| *Smed-RFT2* | dd_Smed_v6_5546_0_1 | SMED30018709 | SMESG000040831.1 |
| *Smed-RFK* | dd_Smed_v6_1500_0_1 | SMED30004205 | SMESG000032740.1 |
| *Smed-FADS* | dd_Smed_v6_8204_0_1 | SMED30019100 | SMESG000076936.1 |
| *Smed-FMNr* | dd_Smed_v6_1834_0_1 | SMED30032272 | SMESG000023871.1 |

**Table S2: Sequences of primers used to generate RNAi probes**

| Gene | Fw | Rv | Planmine reference |
|---|---|---|---|
| *Smed-RFT1* | gtgttggaggtgaaggacgt | agccaaacgtgaccctgtt | dd_Smed_v6_11730_0_1 |
| *Smed-RFT2* | tatctggctgttgtcggtcg | gcgaaccaaacgtcgtcatt | dd_Smed_v6_5546_0_1 |
| *Smed-RFK* | gcgggcaagtaaagaattgg | atgatttccgcttgttctgg | dd_Smed_v6_1500_0_1 |
| *Smed-FADS* | tgattcagtcggcgtcagaa | ccggccaaccttctgtagtt | dd_Smed_v6_8204_0_1 |
| *Smed-FMNr* | gcattaagattggcaaccgaga | tccatgtgcatccccttcag | dd_Smed_v6_1834_0_1 |
| *Smed-β-catenin* | ggcacttcgtcatgttaccc | cggcatattcaaccatttcc | dd_Smed_v6_4850_0_2 |
| *Smed-chat* | tggctcacccagttagagaa | aacaaaatcaatacacgaaaatcaa | dd_Smed_v6_11968_0_1 |
| *Smed-pc-2* | gcatttgtggtgttggagtg | ccaattatggcgtcgatctc | dd_Smed_v6_1566_0_1 |
| *Smed-th* | cttcagaccatccgggttt | acagggttgccaatctttca | dd_Smed_v6_16581_0_1 |
| *Smed-tbh* | acaagcactgccagcaaatg | tccctcaaaagagtcgctgtc | dd_Smed_v6_42610_0_1 |
| *Smed-tph* | ccggtcgacttcattcattt | aaggcgtgtttcaattcacc | dd_Smed_v6_8392_0_2 |
| *Smed-gad* | gtccgatgcaatgaaagagg | gtttggctttccaaagagagg | dd_Smed_v6_12653_0_1 |
| *Smed-lhx1/5-1* | actaagtgcaatgggagccc | caaatatagcctggccccga | dd_Smed_v6_11521_0_1 |
| *Smed-pitx* | cgtcattctccatcggctca | atgcgtctggttctgtcgtt | dd_Smed_v6_15253_0_1 |

**Table S3: Sequences used to generate FISH probes**

| Gene | Sequence |
|---|---|
| *Smed-pc-2* | GGGCYTGGGCGGGGTGAAGGGAAGAGGCTCAGACGAGCGAGGSRGGCTCCCGGCCG<br>CCATGGCCACGGGATTTGAAGGGTGTTTGAGACGGAAGAGAGGCTATAAKCCAATTGRR<br>AAACAAGGTCGTGTCCCACTGCCAACGATTAATAAAAAAGGAATTCCCGGTCTGGGAGA<br>ATCTGACAACATTAAAGTGGATTTGAAATTACCTACTGATCCTCTATTTTCTAAAGAATGGT<br>ACATCAGAAATACCGGTCAAGCTGATGGAGTGAAAGATCTTGATCTTAATGTATTATCAGC<br>ATGGGCTCAAGGTATAACCGGCAAAGGGGTCACGACTGCAATCATGGATGATGGAATAG<br>ACTATCTTCACCCGGATTTATCAAAAAATTATAACGCAGAAAGCAGTTATGATTTCGGTAG<br>CAATGACCCTTACCCGTTCCCTCGATATACTGACGATTGGTTTAACAGTCACGGAACAAG<br>ATGTGCCGGCGAGATTTCCGCCGAACGTGATAATGGCATTTGTGGTGTTGGAGTGGCAT<br>ACGATTCCAAAATCGCTGGGTTGCGCATGCTAGATCAACCATATATGACAGATCTCATTGA<br>GGCAGCAGCTATGAGTTATGATCGAGACAATATTGACATTTACTCTGCTAGCTGGGGACC<br>CACGGATGATGGAAGAACTGTATATGGGCCGAGAAATGAAACGATGCGCGCTATTGTTG<br>AAGGTGTAAATAAGGGACGCAAAGGCAAAGGAAGCATTTACGTTTGGGCATCTGGTGAC<br>GGTGGAGCAGCTGACGATTGTAATTGCGATGGATATGCTGCCAGCATGTGGACAGTAAG<br>TATTAATTCAGCAACCAATGATGGAGAAACCGCAAGCTATGATGAATCCTGTTCTTCAACA<br>CTTGCGTCGACATTTTCTAATGGTAAGGCATGGTTTAGCCTTGACGCCGGTGTCGGAAC<br>TGTGGATTTATATGGCCAATGTACTCTACATCATTCGGGAACTTCTGCTGCAGCTCCAAAA<br>CAGCTGGAGTGTTTGCTCTTGCTCTTGAAGCCAATTCCAATATTGGCTGGAAAAAATGCA<br>CCATTTACAGTGCTTACGATTAACGAAATCTTTGCATGACAAAATCCACCCATAATTGACC<br>TTTATGGACGGGTTGGAATTCATCTCCTTTTGGAAACGGAATACTCAAGCTGGTGAAGTC<br>CCAATTTTCTAAAAGGGGGGTTGTCCAAACATTTCCTGGTTGCCGAAAAATCACGGGGT<br>TTTCTATCAGGTAAAAGTTTAATTGGAAATAACTGACCGGTAATCCGATTGTTTGCAAGAA<br>TTCCTGACGCGCCCGCTTTTTTTTTATTTATTACCCAAGAAAATACAGTTTTTACCCCCCG<br>GGTCCTTTTTGTTCACCCCCACAATAAAGTGTGGCCCCTGGGCAAAATGAGAGCCCCCC<br>CGGGTATGATTTATTTGCCTATACGGGGGCTTATTTTGTGGTTTTCTCCCCGATCACACCA<br>AAATAGAGGCGAGTGAATCCTCTGCCGCCCCCTCCATATTATATAAGAGGAGGTTGGGTT<br>CTACTCCCGGGCGCAATACGAAAAACTCCCCCCCTGATAACTGGCACAAACCCACTGAC<br>CAAATCTCGGCCCTTTGGCGGCGTACGACGGCGGTGACATGGCTGGA |
| *Smed-chat* | GATCTCCTGCCACCAAAGAAACTAGAATATAATATTGATAGCTTTGCCAGTCAAATGATATC<br>TGAAGCCAAACAAGAAATTAAAAATTTGGCCGATAATTTGGATATTTATATTCTACGATACC<br>AGAAATTTGGTAGAGATTTTCCTAAAACTGTCAACATGAGTCCTGATTCCTTCATCCAACT<br>GGCACTTCAACTCACCCATTACAAGATTCATAAATGTCTTGTTCCAACATATGAAAGTGCT<br>TCCATAAGGAGGTTTTCTCTGGGACGCGTAGACAACATTCGTGCGGCTTCAATTCCAGC<br>ATTAAAATGGTGTACATCAATGACTGCTGATACTCCTGCTTCGGATGACGAAAAGATCATT<br>CTTTTACGCTCGGCAATGGAATGGCAGACAAAGGTTATGTTGGACACGATTCAGGGTCA<br>TGGAGTT |
| *Smed-tph* | CGTCGACTGCAGAAAAGACGAGATCAAAAGCTTGATAGAACGTTTGCAACAGAAAATAG<br>ATGACGATTTTCACGTTCAAGAGGTCGACATGATCTCGTCAAATATCAATGGTATTAGAAT<br>TGACACTTTTGATCTAGCAGCGAGCGAGATAGGCGAACAAAACATTCAACTTAAAAACAA<br>AAAACCTGGAATGACAGTGGATAACTTGCCTTGGTTTCCTAAATATATCAAGGATCTAGAC<br>AAAATATCAAATCGCGTTATTATGTATGGATCTGAACTTGATGCTGATCATCCTGGATTTAA<br>AGATGAAGTTTATAGGTTACGTCGAAATCAGTTTTCCGAAATAGCTTACTCATACAAACAT<br>GGAAACCCAATTCCGAAAATTGAATATACAAACGAAGAAATTAAGACTTGGGGAAATGTTT<br>ACCGAGAGTTGACTAAATTATATCCAACTCATGCTTGTCGGGAGTTCTTGCAGAATTTGC<br>CGCTTTTACAAAAGTACTGTGGATACAGAGAAGATAATATTCCTCAACTCGAAGATGTTTC<br>TCAATTTTTAAAGAGTCGATCCGGATTTACTCTACGACCAGTGGCAGGCTATTTGAGTTCT<br>AGAGACTTTTTAGCTGGTCTTGCATTCCGTGTATTTCACTGCACGCAATATATACGCCATC<br>CTTCGGATCCATTTTATACCCCAGAACCGGATTGCTGTCATGAATTAATGGGGCATGTAC<br>CTTTGTTGGCGGATTCAAGTTTTGCTCAGTTTTCTCAAGAAATTGGGTTAGCATCTTTAG<br>GAGCTTCCGAGGACGAAGTTCAAAGATTGGCTACGTGCTACTTCTTCACTATTGAATTTG<br>GACTTTGTCATCAAGATGGAGAACTTCGAGCTTATGGAGCTGGTTTGTTGTCATCTATCG<br>GTGAATTGAAACACGCCTTGAGCAATGAATCTAACATTCAAAAGTTTGACCCAAAAACCG<br>TGATGGAGCAAGAATGCCTAGTGACGACATTTCAGAATGCCTATTTTTTCACTACTTCTTT<br>TGAGGACGCTAAAGATAAAATGAGGGAATTCGCAAAGACAATCAAGAAACCATTCGATGT<br>CCATTACAACCCTTACCCTCAAATGATCGAAACCCTCGACTCCCCAGATTCC |

**Table S4: Sequences of qPCR primers**

| Gene | Fw | Rv | Planmine reference |
|---|---|---|---|
| *Smed-RFT1* | tggaggtgaaggacgttgtg | cctggagccgggtact | dd_Smed_v6_11730_0_1 |
| *Smed-RFT2* | tgtcagttctgccacaatca | tgcactgaaaactggcaa | dd_Smed_v6_5546_0_1 |
| *Smed-RFK* | gcatttgggcggggaagtaa | cccagccgtaataaacccca | dd_Smed_v6_1500_0_1 |
| *Smed-FADS* | acagcttagtaccacatgataaagt | acagtcctttccaccagtga | dd_Smed_v6_8204_0_1 |
| *Smed-FMNr* | cggtgttcctccagctcttt | cccacctgaccagcagaat | dd_Smed_v6_1834_0_1 |
| *Smed-gapdh* | gcaaaacattattccggcttc | gcactggaactctaaaggcca | dd_Smed_v6_78_0_1 |
| *Smed-cys* | aactccatggctagaaccgaa | ccgtcgggtaatccaagtaca | dd_Smed_v6_456_0_1 |
| *Smed-gusb* | gcaaaccagaaacggaaca | ttgtctggataaagggcaca | dd_Smed_v6_3209_0_1 |
| *Smed-hmbs* | tcttgccaaagcaggatt | ccaacggcataaagagaacc | dd_Smed_v6_1346_0_1 |
| *Smed-bactin* | agaacagcttcagcctcgtca | tggaatagtgcttctgggcat | dd_Smed_v6_3_0_1 |
